# Replication fork plasticity upon replication stress requires rapid nuclear actin polymerization

**DOI:** 10.1101/2023.03.24.534097

**Authors:** Maria Dilia Palumbieri, Chiara Merigliano, Daniel González Acosta, Thomas von Känel, Bettina Welter, Henriette Stoy, Jana Krietsch, Svenja Ulferts, Andrea Sanchi, Robert Grosse, Irene Chiolo, Massimo Lopes

## Abstract

Cells rapidly respond to replication stress actively slowing fork progression and inducing fork reversal. How replication fork plasticity is achieved in the context of nuclear organization is currently unknown. Using nuclear actin probes in living and fixed cells, we visualized nuclear actin filaments in unperturbed S phase, rapidly extending in number and thickness upon genotoxic treatments, and taking frequent contact with replication factories. Chemically or genetically impairing nuclear actin polymerization shortly before these treatments prevents active fork slowing and abolishes fork reversal. Defective fork plasticity is linked to reduced recruitment of RAD51 and SMARCAL1 to nascent DNA. Conversely, PRIMPOL gains access to replicating chromatin, promoting unrestrained and discontinuous DNA synthesis, which is associated with increased chromosomal instability and decreased cellular resistance to replication stress. Hence, nuclear F-actin orchestrates replication fork plasticity and is a key molecular determinant in the rapid cellular response to genotoxic treatments.

## Introduction

Interference with the DNA replication process (i.e. replication stress, RS) can be induced by numerous endogenous and exogenous sources^1^ and has recently emerged as a key molecular determinant of genomic instability in early tumorigenesis^2^. Moreover, as tumor cells experience high endogenous levels of RS, additional replication interference by genotoxic treatments or inactivation of key players of the RS response represent promising strategies for cancer chemotherapy^3,4^. Although the RS response is frequently studied upon conditions of severe fork stalling – e.g. by nucleotide depletion or extensive DNA damage – it is crucial to unravel the specific responses to mild replication interference to investigate tumorigenesis and improve therapeutic perspectives, as these conditions are more likely to reflect clinically relevant RS levels.

A key emerging aspect of the RS response in human cells is the plasticity of replication fork architecture^5^. This entails complex unwinding and annealing reactions of DNA strands at replication forks challenged by DNA lesions or other RS sources, and frequently leads to their conversion into 4-way junctions, so-called reversed forks^6^. This transaction promotes an active slowdown of replication fork progression and allows different DNA damage tolerance mechanisms, which overall stabilize stalled forks and promote cellular resistance to genotoxic treatments^5–7^. Several specialized factors were shown to mediate this transaction, including the DNA translocases SMARCAL1, ZRANB3 and HLTF^8–10^, and the central recombinase RAD51^11^. However, reversed forks are also intrinsically unstable intermediates and in certain genetic backgrounds may trigger unscheduled nucleolytic degradation of nascent DNA, contributing to chemosensitivity^12–14^.

An alternative mechanism of replication fork plasticity is provided by the specialized primase PRIMPOL, mediating repriming of DNA synthesis in face of obstacles^5,15^. This discontinuous mode of replication promotes bypass of bulky DNA lesions and implies generation of ssDNA gaps that are filled post-replicatively to complete genome duplication^16–20^. Fork reversal and repriming are competing options of DNA damage tolerance^5^, and fine-tuning their balance recently proved crucial to determine the response to genotoxic treatments^10,21^, and to enable proliferation bursts upon tissue-specific stimuli^22^. Recent evidence suggested that fork plasticity transactions are not limited to forks directly challenged by DNA lesions or replication interference, but rather rapidly extend to unchallenged forks as a global, nuclear response^23^. Although ATR - the central kinase of the human RS response - was implicated in this “signaling” mechanism controlling global fork progression and remodeling^23^, the underlying molecular mechanisms remain elusive and may involve nuclear architecture and dynamics^24^.

Actin is a well-characterized component of the cytoskeleton, involved in multiple cytoplasmic functions, mainly related to its ability to polymerize into filaments (F-actin). Although only a minority of actin resides in the nucleus, nuclear monomeric actin – along with several actin-binding proteins – is a stable component of several chromatin remodeling factors and RNA polymerases^25,26^. These findings provided possible explanations for the relevance of actin in DNA metabolism^27,28^. Filamentous actin structures on the other hand were long undetectable in the nucleus of most cell types. However, recent technological developments in actin filament detection^29^ – i.e. mainly the use of fluorescently labelled actin-binding domains fused to an NLS – allowed detection of nuclear F-actin structures and linked them to various aspects of cell signaling, chromatin dynamics and DNA repair^26,30^. Dynamic and transient F-actin structures were reported in response to serum stimulation^31^ or upon integrin signaling during cell adhesion and spreading^32^. Nuclear F-actin structures are also induced upon T cell receptor (TCR) activation and nuclear accumulation of Ca^2+^, where they appear to control chromatin dynamics and transcription^33,34^. A role for nuclear F-actin in modulating chromatin condensation and nuclear volume was also reported upon mitotic exit^35^.

Importantly, recent evidence has linked nuclear F-actin to different aspects of genome maintenance. Nuclear F-actin structures with different morphologies were described in response to DNA damage^36^. ARP2/3-dependent long and dynamic nuclear actin filaments form in response to ionizing radiation and promote the progression of homologous recombination (HR) repair of heterochromatic double strand breaks (DSB) in *Drosophila*^30,37^ and mouse cells^37^, through myosin-driven directed movement of repair sites to the nuclear periphery. Similarly, ARP2/3-mediated short F-actin filaments clusters DSB to favor their repair by homologous recombination in human cells^30,38,39^. Further, nuclear actin regulates DNA replication initiation upon S-phase entry^40^. A direct contact between the ARP2/3 activator WASP and Replication Protein A (RPA) was recently reported to mediate DNA damage signaling and repair in both human and yeast cells^41^. Moreover, thick, long and persistent nuclear actin filaments were reported in cells experiencing prolonged fork stalling, and were suggested to mediate mobility and repair of broken forks^42^. However, whether nuclear actin polymerization participates in the immediate response to mild replication interference, modulating replication fork progression and plasticity remains elusive.

Here we show that transient nuclear actin filaments are detectable in unperturbed S phase, are rapidly induced by mild genotoxic treatments and are associated with dynamic replication factories. Impairing ARP2/3-mediated nuclear branched actin polymerization by chemical or genetic tools rapidly abolishes active replication fork slowing and remodeling into reversed forks. Moreover, defective nuclear actin polymerization favors the engagement of PRIMPOL over fork remodelers at sites of DNA synthesis, promoting fast and discontinuous DNA synthesis, and impairing chromosome integrity and cellular resistance upon genotoxic treatments. These findings establish transient nuclear F-actin structures as key players of the RS response, paving the way for mechanistic investigations on the role of these and other nuclear architecture components in controlling fork plasticity and the response to chemotherapeutic treatments.

## Results

Distinct and transient nuclear actin filaments form in unperturbed S phase and are induced by replication stress. In order to investigate nuclear actin organization in replicating cells, we imaged stable U2OS cell lines stably expressing the nuclear F-actin marker NLS-actin-chromobody (nAC-GFP) and transiently transfected with the replication fork marker PCNA chromobody (PCNA-CB-RFP) (Fig. 1a). Using our imaging setup, we were able to monitor previously characterized nuclear actin filaments and patches forming upon mitotic exit^35^, as well as short actin structures rapidly induced by treatment with the calcium ionophore A23187^33^ (Fig. 1b; Extended Data Fig. 1a-b; Extended Data Movies 1-2). Intriguingly, unperturbed replicating (PCNA+) cells also display distinct actin filaments, which appear longer and less abundant than those detected upon mitotic exit and after calcium ionophore treatment in similar experimental conditions (Fig. 1b-d; Extended Data Movie 3). Most of the detectable structures are 1-5 μm long and remarkably transient, frequently visible only for a single 20-sec imaging timepoint (Extended Data Fig. 1c-d). To study the impact of mild replication stress on these structures, while imaging the cells, we supplied them with 200nM of the topoisomerase II inhibitor etoposide (ETP), which was previously shown to transiently interfere with DNA replication^11^. ETP treatment induces the formation of new nuclear actin filaments, increasing their overall number in replicating cells, especially within the first imaging period after treatment (Fig. 1d-e; Extended Data Fig. 1e). Hence, distinct transient nuclear F-actin structures are detected in replicating cells and are rapidly induced by mild genotoxic treatments that affect replication fork progression.

**Figure 1.**
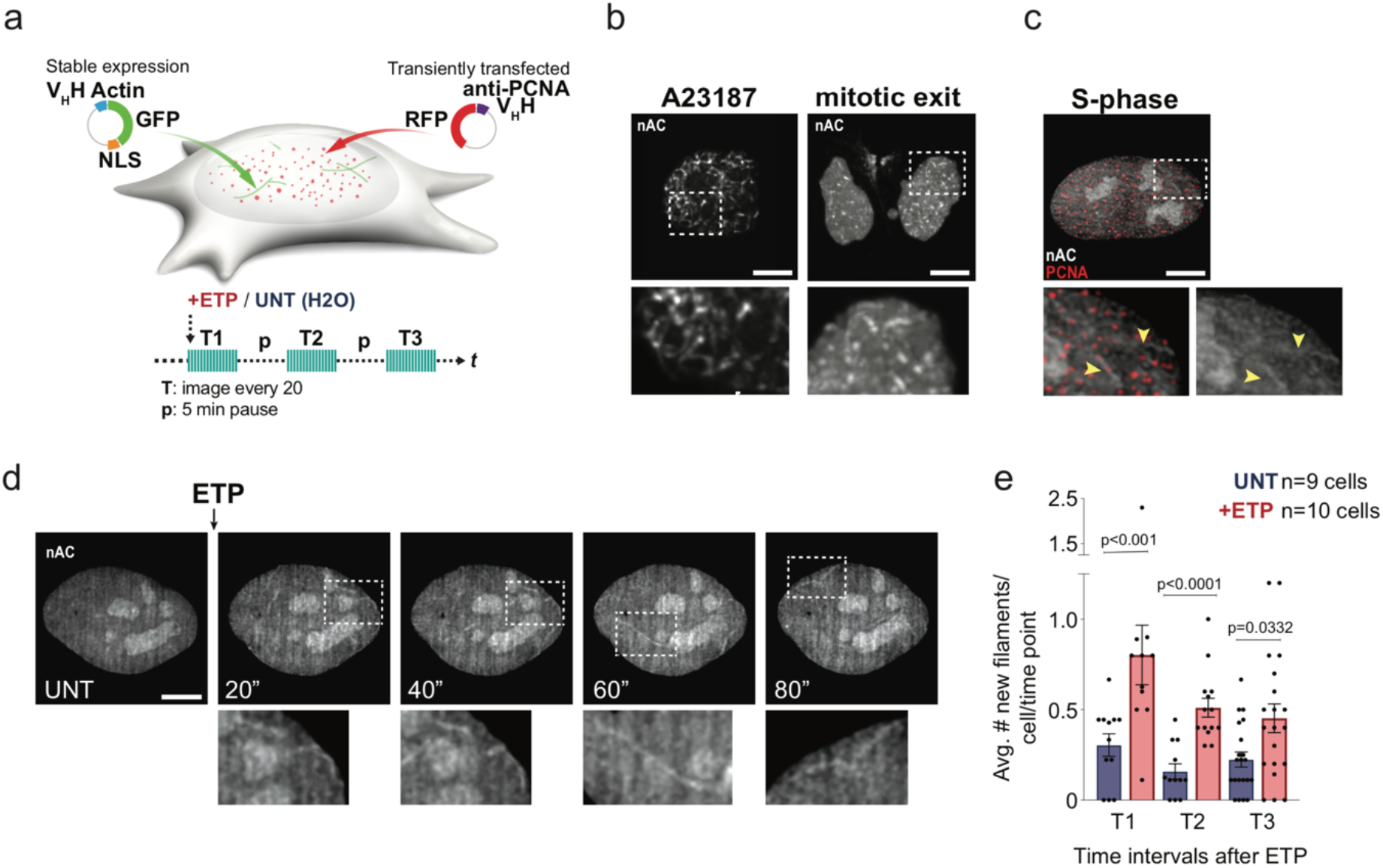
Nuclear actin polymerizes in replicating cells, especially upon mild genotoxic treatments. **a**. Top: Live cell imaging experimental setup: U2OS cells stably expressing nAC-GFP (nuclear actin-chromobody-GFP) were transiently transfected with PCNA-CB-RFP (PCNA-chromobody-RFP) to monitor actin dynamics in replicating cells. Bottom: experimental scheme of the time-lapse imaging experiment in untreated cells (UNT: H2O) or cells treated with 200nM Etoposide (+ETP). Images were taken every 20 seconds for three time intervals (T1,T2,T3) separated by 5 min dark intervals with a spinning-disk confocal microscope. **b**. Representative images from spinning-disk confocal live microscopy analysis of U2OS cells stably expressing nAC-GFP. Cells were imaged after treatment with calcium ionophore A23187 (750 nM) or during mitotic exit, as indicated. See also Extended Data Fig. 1A-B and Extended Data Movies 1-2. **c**. Representative image from spinning-disk confocal live microscopy analysis of S-phase U20S cell (transiently transfected with PCNA-CB-RFP) stably expressing nAC-GFP. Zoomed details highlight examples of transient nuclear actin filaments in PCNA+ cells (yellow arrow in the zoomed detail). **d**. Representative images from spinning-disk confocal live microscopy analysis of PCNA+ U20S cell stably expressing nAC-GFP at indicated time points before (UNT) or after 200 nM ETP treatment. Zoomed details highlight examples of transient nuclear actin filaments. See also Extended Data Movie 3. **e**. Quantification shows the average number of new actin filaments forming in UNT or ETP-treated cells for each cell and time point, during each of the three time intervals described in A. Graph-bars are mean +/-S.E.M. Statistical analysis: numerical p-values for the indicated comparisons are displayed in the figure and calculated with two-tailed Mann–Whitney test. n=9 for the UNT and n=10 for +ETP from 3 or more independent experiments. Images are max intensity projections across a few Z-stacks spanning the nucleus. Scale bars=5 μm.

### Nuclear actin filaments contact replication factories and correlate with their increased mobility

Although live-cell imaging by actin chromobody is a powerful tool to investigate the dynamics of nuclear F-actin ^33,35,42^, it is intrinsically limited in its detection power by relatively low signal/noise ratio and may visualize only a subset of particularly visible structures^29,37,42^. To complement these observations, we investigated nuclear F-actin by immunofluorescence analysis of U20S cells transiently expressing FLAG-NLS-Actin^43,44^, where replication factories are identified by EdU incorporation. Fixed cell imaging reveals a dense network of nuclear F-actin structures, including fine and branched filaments (thin filaments), bright punctate structures (thick short filaments) and bright elongated structures (thick long filaments (Fig. 2a). 85% of F-actin-positive cells are also EdU-positive, indicating that filaments are mostly associated with replicating cells. Detection of these structures in the middle Z-stacks of confocal images confirms that they are nuclear (Extended Data Fig. 2a). Both filamentous and punctate signals reflect polymerized actin, as they are largely lost in cells expressing the actin mutant R62D (FLAG-NLS-R62D-Actin), which acts as a dominant negative poisoning actin polymerization^35^ (Extended data Fig. 2b). We then treated cells with ETP or with mild doses of the topoisomerase I inhibitor camptothecin (CPT), also previously shown to affect replication fork structure and architecture with marginal effects on chromosome integrity and cell survival^11,45^. These treatments result in an increased number of cells displaying short and long thick filaments, suggesting actin polymerization or bundling in response to replication stress (Fig. 2a-b; Extended Data Fig. 2c). Most thick short filaments are proximal or overlapping with EdU foci (Fig. 2c), consistent with extensive actin polymerization occurring at replication sites.

**Figure 2.**
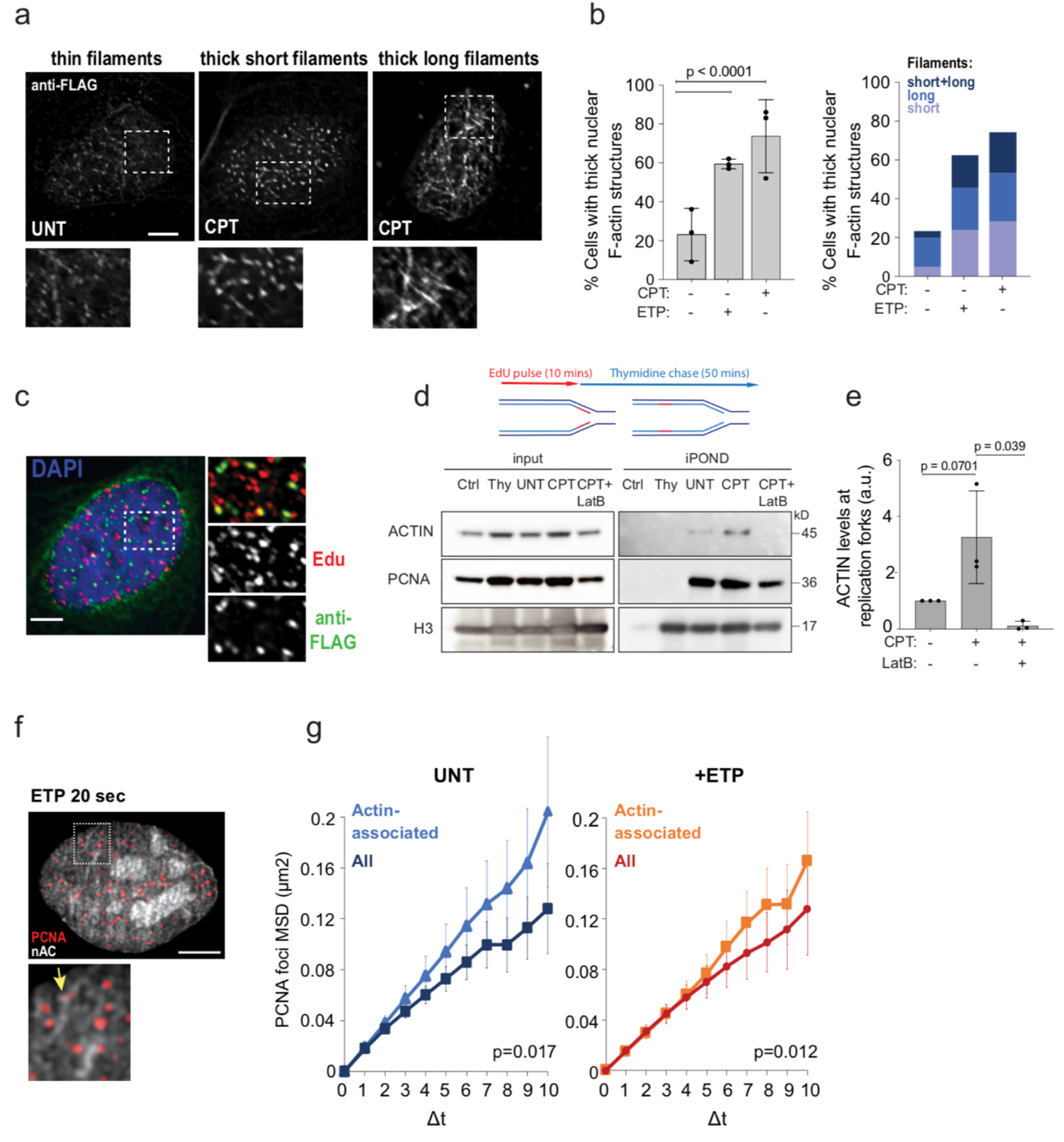
Nuclear F-actin interacts with the replication factories and locally affects their dynamics. **a**. Deconvolution microscopy image of fixed S-phase (EdU+) U2OS cells transiently transfected with FLAG-NLS-WT-Actin, and stained for FLAG and EdU by immunofluorescence (IF). Cells are untreated (UNT) or treated with 100 nM camptothecin (+CPT) (or 200 nM etoposide (+ETP) in Extended Data Fig. 2c) for 1 hour and subjected to EdU incorporation for 4 minutes before fixation. Images show thin filaments in UNT and short and long thick filaments in +CPT cells. Zoomed details highlight examples of the actin filaments. **b**. Quantifications of the experiment described in A. Data are mean +/-SEM; N= 91, 58 and 60 cells for UNT, +CTP and +ETP respectively, from three independent experiments. Statistical analysis: p-values for the indicated comparisons were calculated with two-tailed Mann-Whitney test. **c**. IF show examples of short actin filaments colocalizing with EdU in a CPT treated cell (+CTP). **d**. Isolation of proteins on nascent DNA (iPOND). Top: HEK293T cells were treated and EdU-labeled as indicated in Extended Data Fig. 2d. 10 µM thymidine (Thy) was added for 50 minutes after the 10 minutes EdU labeling and is used to discriminate proteins associated with chromatin behind replicating forks. Bottom: Western blots of input and iPOND purified proteins from indicated treatments. Cells were treated with 100 nM CPT for 1 hour. 100 nM LatB was optionally added 10 minutes before CPT and retained during the genotoxic treatment (See Extended Data 2D). Proteins associated with nascent DNA were isolated by iPOND and detected with the indicated antibodies. In the control (Ctrl) experiment, the click reaction is performed using DMSO instead of biotin azide. **e**. Graph-bar depicts mean and SD of quantified ACTIN levels at nascent DNA from three independent iPOND experiments. Values were normalized to the H3 control and are represented as fold change over the UNT sample (a.u. = arbitrary units). Statistical analysis: numerical p-values for the indicated comparisons were calculated with one-tailed t-test with Welch’s correction. **f**. Representative image of a cell from the experiment described in Fig. 1a shows an example of PCNA focus colocalizing with an actin filament (yellow arrow in the zoomed detail). **g**. MSD analysis of nuclear F-actin-associated PCNA foci relative to all PCNA foci in UNT or +ETP cells. n=6 cells for the actin-associated PCNA foci and n=9 cells for All PCNA foci in UNT. n=6 cells for the actin-associated PCNA foci and n=8 cells for the All PCNA foci in +ETP cells, from 3 or more independent experiments. Data are mean +/-S.E.M. p-values were calculated with extra sum-of-squares *F*-test, nonlinear regression for curve fitting. Δ*t*, time intervals (intervals were 20 s each). Scale bars=5 μm.

We further assessed the association of nuclear F-actin and DNA replication centers by isolation of proteins on nascent DNA (iPOND^46^) (Fig. 2d-e; Extended Data Fig. 2d). Immunoprecipitation of nascent DNA identifies actin at DNA synthesis centers, confirming that actin is detected proximal to replication factories in unperturbed S phase (Fig. 2d). Remarkably, this interaction is significantly enhanced upon CPT treatment and lost when the actin polymerization inhibitor LatrunculinB (LatB) is added to the media shortly (10 min) before CPT (Fig. 2d-e; Extended Data Fig. 2d). We then took advantage of our live-cell imaging setup to investigate the impact of nuclear F-actin on the dynamics of replication factories, monitoring their mobility over time via mean square displacement (MSD) analysis. While ETP treatment does not result in a detectable increase of MSD values for the bulk population of PCNA foci (Extended Data Fig. 2e), F-actin-associated PCNA foci (Fig. 2f) display significantly higher mobility, both in untreated and ETP-treated cells (Fig. 2g). Overall, we conclude that nuclear F-actin establishes detectable contacts with replication factories and affects their dynamics.

### ARP2/3-dependent nuclear actin polymerisation is required for active fork slowing upon replication stress

We next investigated the functional relevance of the interaction between nuclear F-actin and replication factories on DNA replication, in presence or absence of mild genotoxic treatments. To this purpose, we analysed replication fork progression at single-molecule level by spread DNA fiber assays^47^, providing cells with halogenated nucleotides and mild doses of ETP or CPT. These treatments were previously shown to induce marked fork slowing and reversal, with no major impact on cell cycle progression and cell viability^11^. In this setup, we induced actin depolymerization - adding LatB or Swinholide A (Swi), an alternative actin depolymerization agent - when incorporation of halogenated nucleotides was already ongoing, i.e. 10 min before addition of the genotoxic drugs (Fig. 3a). Pre-treatment with LatB or Swi does not detectably affect replication fork progression (Extended Data Fig. 3a), showing that actin polymerization is *per se* not required to support efficient fork progression in unperturbed conditions. As expected from previous studies, both ETP and CPT drastically affects replication fork progression, but the active fork slowing observed in these conditions is significantly rescued by either of the actin polymerization inhibitors (Fig. 3a-c and Extended Data Fig. 3b-c). Although actin polymerization can be promoted by various co-factors, branched actin filament formation uniquely requires the ARP2/3 complex, which can be specifically inhibited by CK666 or CK869 treatment^48^. Similarly to LatB or Swi, 10 min pretreatment with either ARP2/3-inhibitor has no impact on unperturbed DNA synthesis *per se* (Extended Data Fig. 3a), but abolishes CPT-induced fork slowing (Fig. 3d and Extended Data Fig. 3d). Moreover, siRNA-mediated downregulation of Arp3 has very similar effects (Extended Data Fig. 3e-g), suggesting that the branched actin network plays a pivotal role in modulating replication fork progression upon DNA damage.

**Figure 3.**
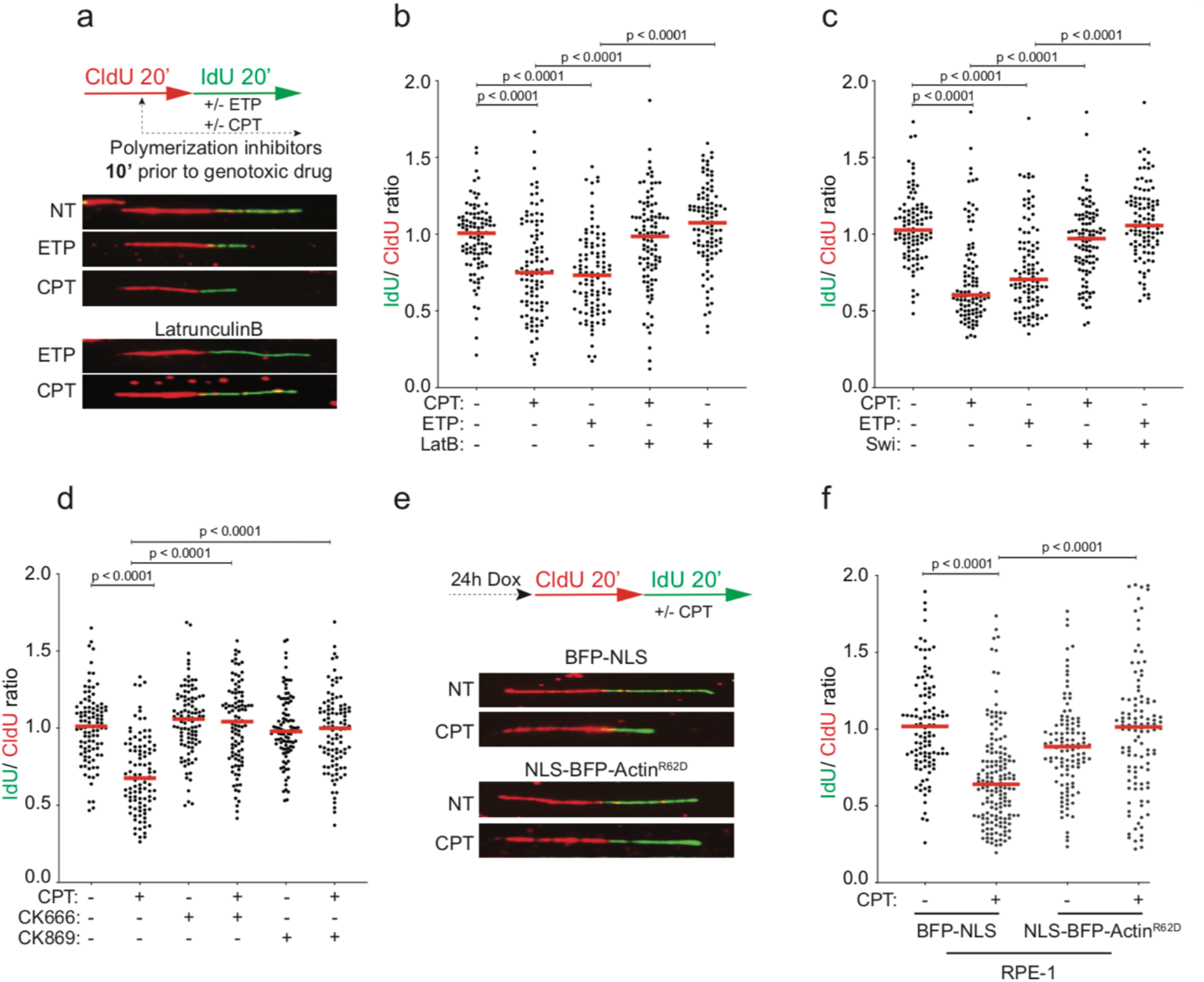
Nuclear actin polymerization is required for active fork slowing upon mild genotoxic stress. **a**. DNA fiber analysis of U2OS cells. Top: schematic of the CldU/IdU pulse-labeling protocol used to evaluate fork progression upon 100 nM CPT and 20 nM ETP. 100 nM LatB, Swi, CK666 or CK869 were added 10 minutes prior to CPT or ETP and retained during the IdU labelling. Bottom: representative DNA fibers. **b-d**, f.The IdU/CIdU ratio is plotted as a readout of fork progression. A minimum of 100 forks (indicated as black dots) was scored in three independent experiments yielding similar results. The median values are indicated by horizontal red lines. See Extended Data Fig. 3b-d,h for compiled repetitions. Statistical analysis: numerical p-values for the indicated comparisons are displayed in the figure and calculated with Mann–Whitney test. **e-f**. DNA fiber analysis of RPE-1 cells stably expressing doxycycline inducible BFP-NLS or NLS-BFP-ActinR62D. Top: schematic of the CldU/IdU pulse-labeling protocol used to evaluate fork progression upon 100 nM CPT. Doxyclicline (Dox) was added 24 hours before CldU/IdU pulse-labeling. Left: representative DNA fibers are shown for each condition.

Although the minimized timing of treatment excludes long-term pleiotropic effects, chemical inhibition of actin polymerization may unavoidably affect the bulk actin network in the cytoplasm. To specifically investigate the functional relevance of nuclear F-actin in the modulation of replication fork progression, we took advantage of a previously established genetic tool, i.e. a stable RPE-1 cell line bearing a doxycycline-inducible R62D actin mutant (NLS-BFP-Actin^R62D^)^32,35,43^. Thanks to the NLS, this exogenous actin specifically impairs nuclear actin polymerisation, but does not detectably affect cytoplasmic F-actin and its functions^35,37^. Remarkably, induction of NLS-Actin^R62D^ 24h before our fiber assays also fully suppresses CPT-induced fork slowing (Fig. 3e-f and Extended Data Fig. 3h), establishing nuclear F-actin as key molecular determinant of the rapid modulation of replication fork progression upon mild genotoxic treatments.

### Nuclear actin polymerization mediates the engagement of fork remodelling factors and fork reversal

Active replication fork slowing upon DNA damage or mild replication interference was recently linked to replication fork reversal, i.e. the controlled and reversible remodelling of replication forks into four-way junctions^5^. This transaction requires the recruitment to forks of the RAD51 recombinase^11^ and active engagement of the specialized translocase SMARCAL1, which is a stable component of the replisome^8,49^. We thus used the iPOND approach described in Fig. 2 to investigate whether nuclear F-actin may be required to enable recruitment or engagement of these remodelling factors on nascent DNA. In Consistent with previous studies^8^, RAD51 presence on nascent DNA is significantly increased upon CPT treatment, while SMARCAL1 is detectable at replication forks at comparable levels in treated and untreated cells (Fig. 4a-b). Remarkably, treating cells with LatB 10 min prior to CPT markedly reduces the levels of both RAD51 and SMARCAL1 on nascent DNA, suggesting defective recruitment to replication forks or defective engagement of these factors at sites of DNA synthesis, which may prevent efficient crosslinking and detection on nascent DNA.

**Figure 4.**
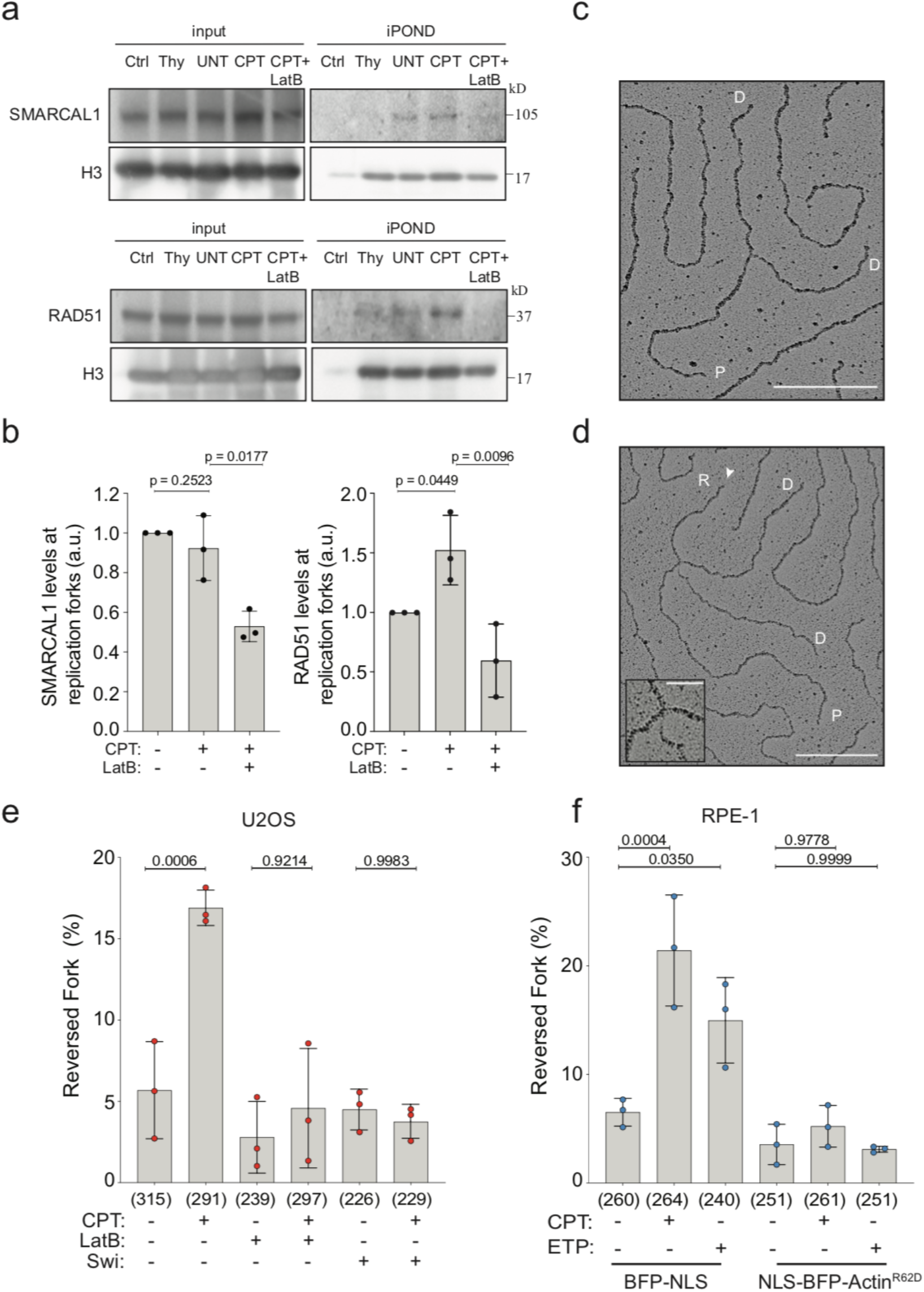
Nuclear F-actin modulates the engagement of replication fork remodellers and fork reversal. **a**. iPOND. HEK293T cells were treated and EdU labeled as indicated in Extended Data 2D. Western blots of input and iPOND purified proteins from indicated treatments. Cells were treated with 100 nM CPT for 1 hour. 100 nM LatB was optionally added 10 minutes before CPT and retained during the genotoxic treatment (See Extended Data 2d). Proteins associated with nascent DNA were isolated by iPOND and detected with the indicated antibodies. In the control (Ctrl) experiment, the click reaction is performed using DMSO instead of biotin azide. **b**. Graph-bar depicts mean and SD of quantified RAD51 and SMARCAL1 levels at nascent DNA from three independent iPOND experiments. Values were normalized to the H3 control and are represented as fold change over the UNT sample (a.u. = arbitrary units). Statistical analysis: numerical p-values for the indicated comparisons were calculated with one-tailed t-test with Welch’s correction. **c-d** Electron micrographs of representative replication forks from U2OS cells: parental (P) and daughter (D) duplexes. d. The white arrow indicates the regressed arm (R); the four-way junction at the reversed fork is magnified in the inset. Scale bar = 200 nm, 40 nm in the inset. **e**. Frequency of reversed replication forks isolated from U2OS cells upon optional treatment with 100 nM CPT for 1 hour. 100nM LatB or Swi were added 10 minutes before CPT and retained during the genotoxic treatment. In brackets, total number of molecules analyzed per condition. **f**. Frequency of reversed replication forks isolated from RPE-1 cells after 24 hours doxycycline-induction of either BFP-NLS or NLS-BFP-ActinR62D and optionally treated for 1 hour with 100 nM CPT or 20 nM ETP. In brackets, total number of molecules analyzed per condition. **e-f**.The graphs depict mean and SD from three independent EM experiments. Statistical analysis: numerical p-values for the indicated comparisons are displayed in the figure and calculated with ordinary one-way ANOVA.

We then directly assessed whether defective active polymerization affects the ability of the cells to promptly induce replication fork reversal upon genotoxic stress. For this purpose, we took advantage of an established approach for direct electron microscopic visualization of replication intermediates^50,51^, which allows distinguishing standard 3-way replication forks from 4-way reversed forks (Fig. 4c-d). In line with previously published data^11^, both U2OS and RPE-1 cells display a drastic and reproducible increase in the percentage of reversed forks upon CPT or ETP treatment (Fig. 4e-f and Extended data Fig. 4a-b). Remarkably, treatment with either LatB or Swi 10 min before CPT treatment fully abolishes drug-induced replication fork reversal in U2OS cells (Fig. 4e and Extended data Fig. 4a). Moreover, expression of the NLS-Actin^R62D^ dominan-negative mutant also impaired CPT- and ETP-induced reversal in RPE-1 cells (Fig. 4f and Extended data Fig. 4b), establishing nuclear actin polymerization as a strict requirement for active fork slowing and reversal in human cells.

Previous work had linked the central kinase of the replication stress response ATR with efficient replication fork reversal^23^. Moreover, nuclear F-actin assembly was recently suggested to mediate local ATR activation upon laser microirradiation^52^. However, we do not detect any defects in canonical markers of global ATR activation upon CPT treatment – i.e. ATR autophosphorylation or phosphorylation of the key partner kinase CHK1 (Extended Data Fig. 4c) – suggesting that the role of nuclear F-actin in replication fork progression and remodelling is independent or downstream from canonical ATR activation.

### Defective nuclear actin polymerization promotes PRIMPOL-mediated discontinuous DNA synthesis

We next investigated the genetic dependencies of the unrestrained fork progression observed upon replication interference when nuclear actin polymerization is impaired. Accelerated DNA synthesis was previously reported upon PARP inhibition and was linked to deregulated fork restart activity of the RECQ1 helicase, which prevents fork pausing in the reversed state^11,53,54^. However, effective downregulation of RECQ1 by siRNA does not restore active fork slowing upon ETP treatment, in the presence of LatB (Extended Data Fig. 5a-c). Thus, unrestrained fork progression upon defective nuclear actin polymerization does not reflect accelerated restart of previously reversed forks. We next assessed whether defective fork slowing was linked to deregulated *de novo* restart of DNA synthesis on a damaged template, which was recently reported in other genetic conditions impairing fork reversal^10,21,55^. This discontinuous mode of DNA synthesis implies the transient formation of ssDNA gaps on newly replicated duplexes, and can be detected in a modified DNA fiber assay as shortening of replicated tracks by cleavage of the ssDNA-specific S1 nuclease, prior to fiber stretching on microscopy slides^56^ (Fig. 5a). Indeed, S1-induced replication track shortening is specifically detected in our assays when ETP was combined with LatB pretreatment (Fig. 5b and Extended data Fig. 5d), confirming that unrestrained fork progression upon defective nuclear actin polymerization entails discontinuous DNA synthesis on a damaged template.

**Figure 5.**
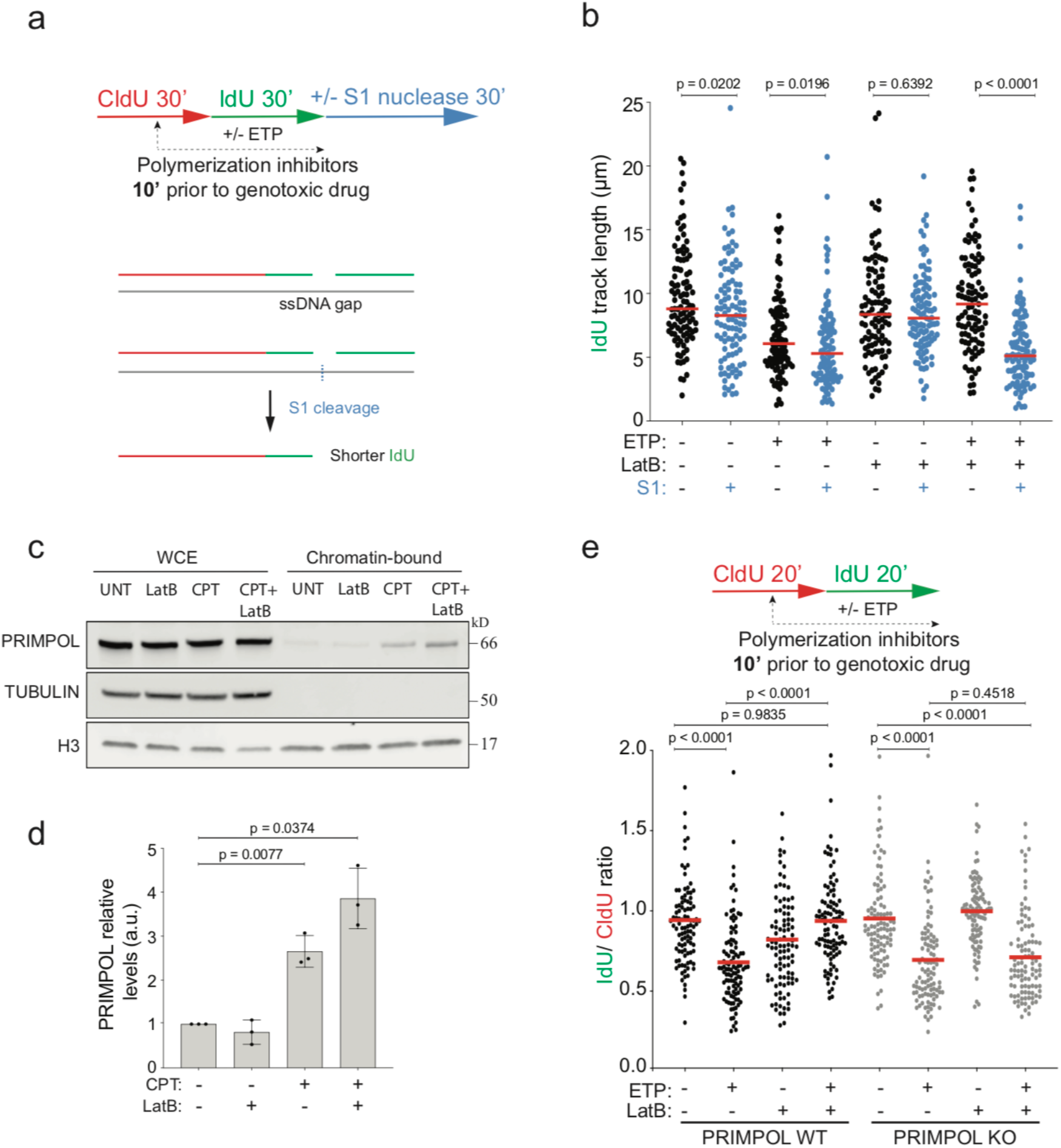
Defective actin polymerization promotes PRIMPOL-mediated fast and discontinuous DNA synthesis. **a**. Schematic of the CldU/IdU pulse-labeling protocol coupled with 30 minutes ssDNA-specific S1 nuclease treatment for the detection of ssDNA gaps on ongoing forks upon optional treatment with 20 nM ETP. 100 nM LatB was added 10 minutes prior to ETP and retained during the IdU labelling. **B**. The IdU track length (μm) is plotted as a readout of discontinuous DNA synthesis. A minimum of 100 forks was scored in three independent experiments yielding similar results. The median values are indicated by horizontal red lines. See Extended Data Fig. 5d for compiled repetitions. Statistical analysis: numerical P-values for the indicated comparisons were calculated with Mann–Whitney test. **c**. Immunoblot detection of the indicated proteins in whole cell extracts (WCE) or chromatin bound fraction. **d**. Graph-bar depicts mean and SD of quantified PRIMPOL levels in the chromatin bound fraction from three independent chromatin fractionation experiments. Values were normalized to the H3 control and are represented as fold change over the UNT sample (a.u. = arbitrary units). Statistical analysis: Numerical p-values for the indicated comparisons are displayed in the figure and calculated with one-tailed t-test with Welch’s correction. **e**. DNA fiber analysis of U2OS PRIMPOL WT and KO cells. Top: schematic of the CldU/IdU pulse-labeling protocol used to evaluate fork progression upon 20 nM ETP. 100 nM LatB was added 10 minutes prior to ETP and retained during the IdU labelling. Bottom: the IdU/CIdU ratio is plotted as a readout of fork progression. A minimum of 100 forks was scored in three independent experiments yielding similar results. The median values are indicated by horizontal red lines. See Extended Data Fig. 5e for compiled repetitions. Statistical analysis: numerical P-values for the indicated comparisons were calculated with Mann–Whitney test.

As this discontinuous DNA synthesis was previously linked to the action of the PRIMPOL primase^10,15–18,21^, we investigated PRIMPOL engagement in DNA synthesis in our experimental conditions. As PRIMPOL could not be detected in iPOND experiments with currently available procedures and reagents (data not shown), we relied on chromatin fractionation experiments and indeed observed that PRIMPOL recruitment to chromatin is induced by CPT treatment – as previously reported upon induction of DNA damage^16,19,55^ – and further enhanced by pre-treatment with LatB (Fig. 5c-d). We next directly assessed the genetic contribution of PRIMPOL performing DNA fiber assays on PRIMPOL-KO U2OS cells, and confirmed that the unrestrained fork progression induced by LatB in the presence of ETP is indeed entirely dependent on PRIMPOL (Fig. 5e and Extended Data Fig. 5e). Importantly, PRIMPOL dependency for unrestrained fork progression is also observed when ARP2/3-dependent branched nuclear F-actin is impaired by CK666 treatment (Extended Data Fig. 5f-g). Hence, defective nuclear actin polymerization rapidly provides deregulated access of PRIMPOL to damaged replication forks, promoting excessive repriming and discontinuous DNA synthesis.

### Nuclear actin polymerization limits genomic instability and cellular sensitivity upon DNA damage

Finally, we investigated the functional consequences of nuclear F-actin deregulation on the cellular response to replication stress, in terms of genome stability and cell survival. We first used chromosome spreads from metaphase arrested cells to monitor chromosomal breaks and abnormalities (Fig. 6a). Using mild CPT treatments that do not induce *per se* significant chromosomal instability in U2OS cells, we observe a marked increase of chromosomal instability when actin polymerization is impaired by LatB shortly before CPT treatment (Fig. 6b). Similarly, specific impairment of nuclear F-actin by inducible expression of NLS-Actin^R62D^ in RPE-1 cells increases CPT-induced chromosomal abnormalities (Fig. 6c). Moreover, clonogenic assays performed in in NLS-Actin^R62D^ RPE-1 cells show that impairment of nuclear actin polymerization decreases cell survival upon CPT treatment (Fig. 6d-e). Overall, these data establish the functional relevance of rapid nuclear actin polymerization for efficient DNA damage tolerance during replication.

**Figure 6.**
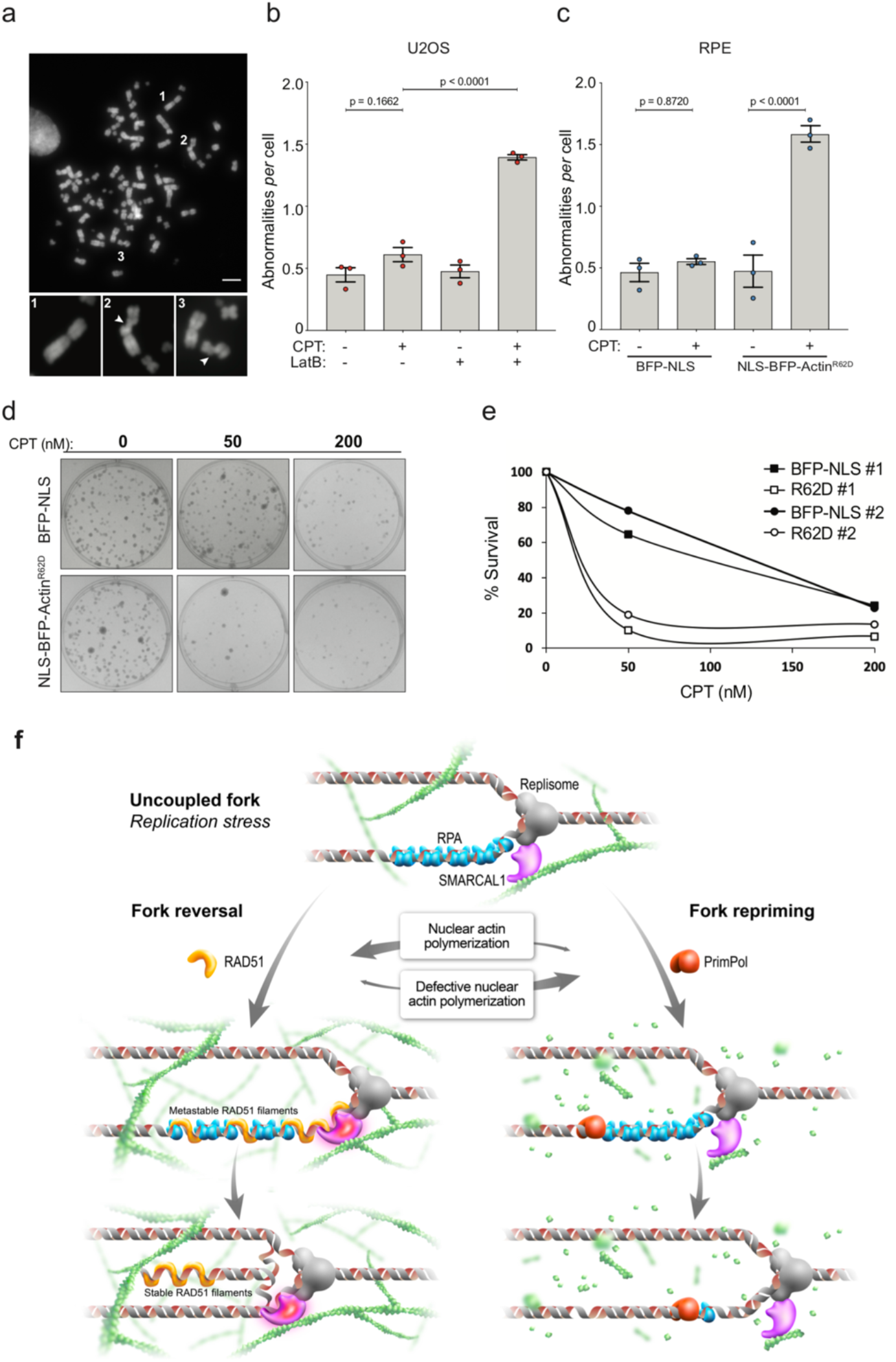
Nuclear F-actin limits chromosomal instability and cell lethality upon genotoxic treatments. **a**. Representative metaphase spread image. Scale-bar=5 μm; 1 representative intact chromosome and 2, 3 representative breaks. **b**. Number of chromosomal abnormalities in U2OS cells optionally treated with 100 nM CPT for 2 hours. 100 nM LatB was added 10 minutes before CPT and retained during the genotoxic treatment. **c**. Number of chromosomal abnormalities in RPE-1 cells stably expressing doxycycline inducible NLS-Actin-WT or NLS-Actin-R62D. Doxycycline (Dox) was added for 24 hours prior to the experiment and cells were optionally treated for 8 hours with 50 nM CPT. **b-c**. The graph-bar depicts mean and SD from three independent experiments. A minimum of 50 metaphases was analyzed per sample. Statistical analysis: numerical P-values for the indicated comparisons were calculated with one-way ordinary ANOVA. d. Representative images of E. **e**. Colony formation assay of RPE-1 cells after 24 hours doxycycline-induction of either BFP-NLS or NLS-BFP-ActinR62D and optionally treatment with 50 nM or 200 nM CPT for 1 hour. Survival is displayed as relative number of cells forming colonies, compared to UNT (set at 100%). Data are represented as mean of 3 measurements for each of the two displayed experiments. **f** Working hypothesis: nuclear F-actin polymerization facilitates RAD51 recruitment at stalled/uncoupled forks, promoting SMARCAL1-dependent fork reversal and limiting PRIMPOL-mediated repriming (left). Defective nuclear actin polymerization leads to deregulated PRIMPOL recruitment, promoting unrestrained, discontinuous DNA synthesis (right; see Discussion for details).

## Discussion

Our data identify a novel role of nuclear F-actin structures in DNA replication in human cells. We provide several lines of evidence that nuclear F-actin plays a pivotal role in orchestrating the rapid response to replication interference, promoting chromosome stability and cell tolerance of genotoxic treatments. Specifically, we detect transient and dynamic F-actin structures that form in a normal S-phase, are rapidly induced by replication stress, and are associated with replication sites. F-actin structures observed in this context are distinct from previous filaments and patches previously detected by actin-CB expression in response to various stimuli^31,32,34,35^ – including prolonged fork stalling^42^ – and may have previously escaped detection by live imaging given their thin structure, transient nature, and low abundance. Importantly, fixed cell imaging studies reveal a complex network of actin filaments establishing frequent contacts with replication sites, particularly in response to replication stress, consistent with a global role of F-actin in replication fork plasticity.

Moreover, we show that nuclear actin polymerization acts as a critical determinant in the choice between alternative mechanisms of DNA damage tolerance during replication, i.e. replication fork reversal vs repriming. Both mechanisms require accumulation of RPA-coated ssDNA^11,16^, which is typically observed upon stalling of leading strand synthesis at DNA lesions, while unwinding by the replicative helicase and lagging strand synthesis proceed beyond the lesion (uncoupled fork; Fig. 6f). However, while PRIMPOL-mediated repriming is promoted by direct interaction with RPA and *de novo* DNA synthesis by its primase activity^57^, replication fork reversal is a complex reaction^5^, requiring partial exchange of RPA with RAD51 – catalysed by the action of RAD51 paralogs^11,58^ – and the concerted action of SMARCAL1, ZRANB3 and HLTF translocases^8–10^, differentially activated by RAD51 and its cofactors^49^ (Fig. 6f). Nuclear F-actin may create a favorable local environment to recruit fork remodellers and/or their activators at uncoupled replication forks, thereby triggering fork reversal and limiting PRIMPOL engagement on RPA-coated DNA. Alternatively, polymerizing branched F-actin in proximity to replication factories may prevent deregulated access of PRIMPOL to ssDNA, which would lead to an excessively discontinuous DNA synthesis and may saturate the cellular capacity for post-replicative gap-filling^18,20^. Considering that RPA-coated ssDNA is proposed to extrude as a loop from the replication center, due to physical interactions between the stalled polymerase and the moving helicase, preventing unlimited access of PRIMPOL to RPA-coated ssDNA may be required to stabilize the uncoupled fork and to kinetically allow the complex biochemical reactions required for fork reversal. In this context, defective nuclear actin polymerization could provide deregulated access of PRIMPOL to ssDNA gaps at uncoupled forks, thereby preventing fork remodeling (Fig. 6f).

Several hypotheses are currently open on how nuclear F-actin may rapidly and globally affect replication fork plasticity upon genotoxic treatments, and will represent fascinating avenues for future research. First, similarly to what described upon mitotic exit^35^, nuclear F-actin may be required upon RS to locally modify chromatin compaction. However, differently from the global chromatin decondensation observed during nuclear volume expansion in G1 cells, replicating cells may use the nuclear F-actin network to induce local changes in chromatin compaction, which could help executing the replication program and responding rapidly to RS. Intriguingly, changes in chromatin compaction were reported upon multiple sources of RS, linked to specific epigenetic marks and proposed to mediate nuclear positioning of forks experiencing prolonged stalling^23^.

Second, nuclear F-actin can affect fork plasticity by modifying the nuclear position or dynamics of replication sites. Differently from DSB repair^30,37,39^ and prolonged fork stalling or collapse^42^, we did not detect global relocation of replication factories within the experimental time frame upon mild RS,, suggesting that extensive relocalization is not needed to promote fork reversal or inhibit PRIMPOL engagement. However, the observed increased mobility of PCNA foci linked to the most visible F-actin structures by live imaging suggests that dynamic movements are part of the replication stress response and may become detectable only in certain contexts, e.g. in the presence of persistent fork stalling or breakage^42^. It is also possible that short-range movement of bulk replication sites – hardly detectable with current imaging resolution - contributes to orchestrate replication fork plasticity. Sudden changes in chromosome dynamics promote homologous pairing in meiosis^59^; by analogy it is possible that similarly complex biochemical reactions – such as extensive DNA unwinding and annealing driving fork reversal – may require increased short-range mobility of nascent DNA, mediated by local nuclear actin polymerization. Finally, as chromosome mobility and directed movement is linked to efficient DNA repair^37,39,60–65^, it will be essential to thoroughly investigate the functional relevance of myosin and other motor proteins in modulating replication fork plasticity, as a rapid response to replication interference. In light of the surprising impact of nuclear F-actin on chromosome integrity and cell survival upon mild replication interference, specific players involved in nuclear actin polymerization may be key determinants of the response to cancer chemotherapy, highlighting the clinical relevance of further mechanistic investigations in this area.

## Main Figures and legends

## Extended Data Figures and legends

**Extended Data Figure 1.**
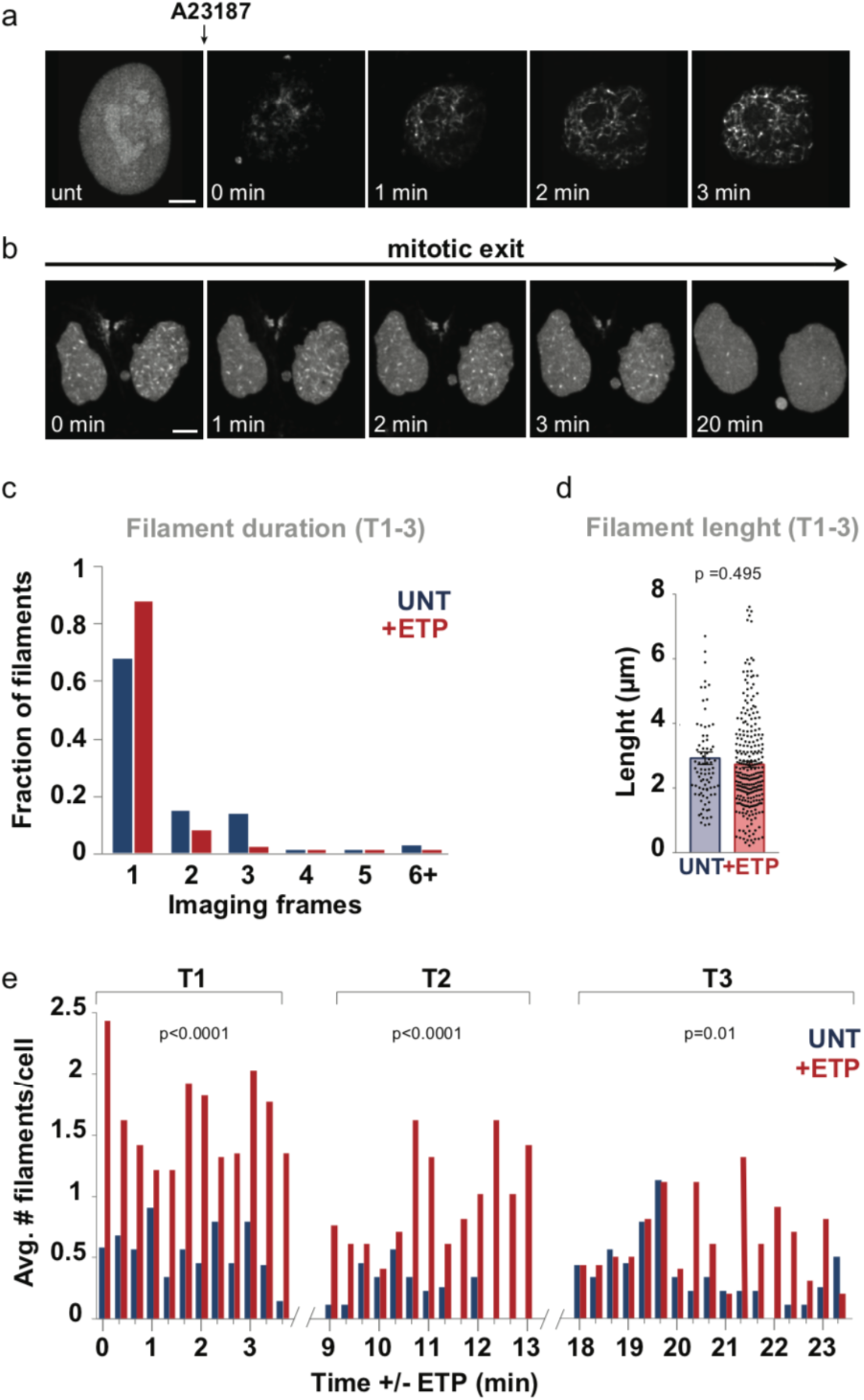
Related to Fig. 1. **a-b**. Snapshots from a time-lapse experiment of U20S cells stably expressing nAC-GFP. Cells were imaged after 750 nM A23187 treatment or during mitotic exit, as indicated. Scale bars=5 μm. c. Duration of filaments *per* imaging frame (20 seconds) in untreated cells (UNT) or treated with 200nM Etoposide (+ETP) from the experiment described in Figure 1a. d. Actin filament length observed in UNT or ETP treated cells from the experiment described in Figure 1A. **e**. Average number of filaments *per* cell/time point (T1,T2,T3) in UNT or ETP treated cells. Data are mean ± S.E.M, N*=*84 filaments (9 cells) for UNT and N*=* 273 filaments (10 cells) for +ETP in **c, d** and **e**. Statistical analysis: numerical P-values for the indicated comparisons were calculated with two-tailed Mann–Whitney test in d and e.

**Extended Data Figure 2.**
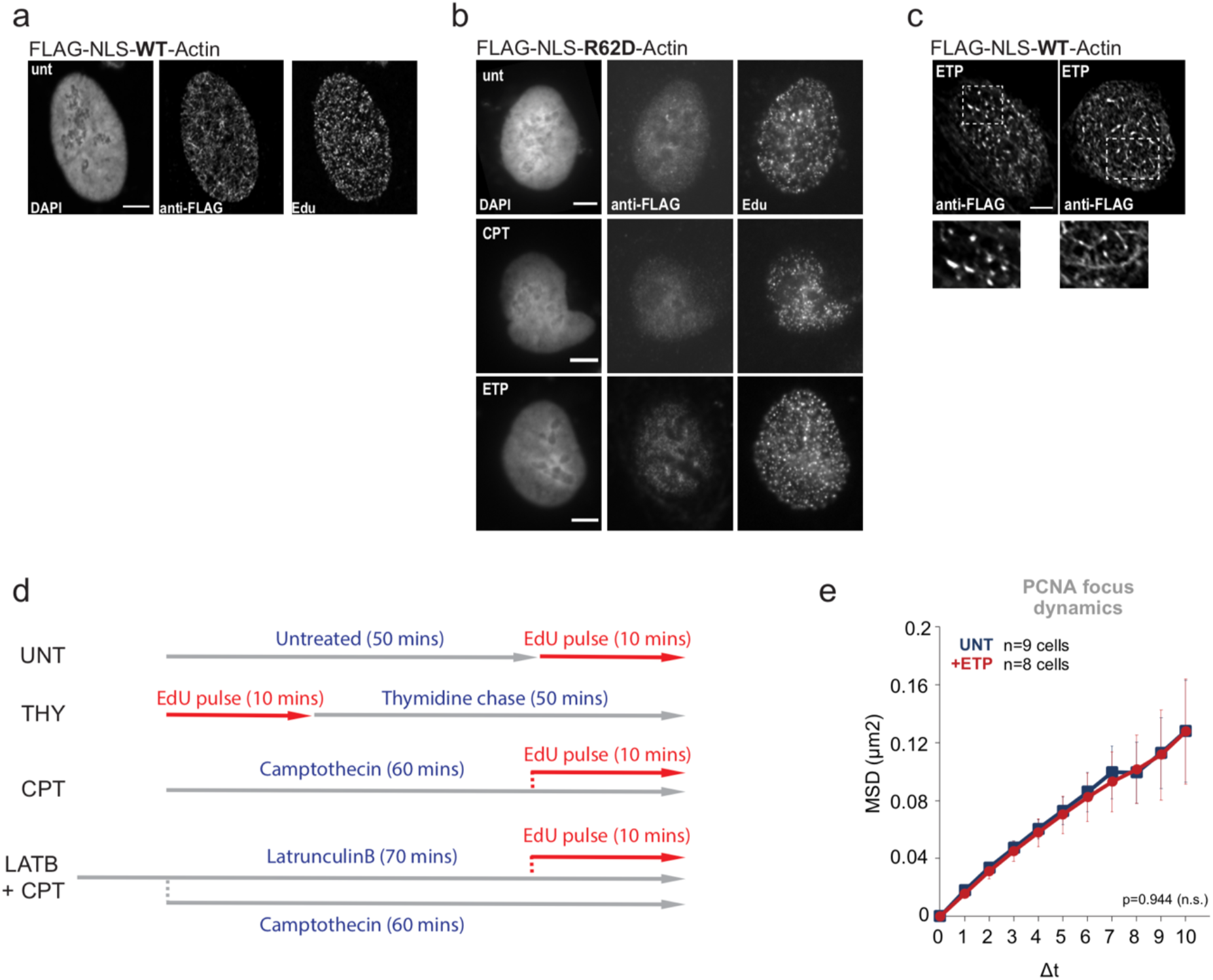
Related to Fig. 2. **a**. Confocal microscopy of a single middle *Z-*stack from fixed S-phase (EdU+) U2OS cell transiently transfected with FLAG-NLS-WT-Actin and stained for FLAG and EdU. b. Representative confocal images of fixed S-phase (EdU+) U2OS cells transiently transfected with FLAG-NLS-R62D-Actin and stained for FLAG and EdU. Cells are untreated (UNT), treated with 100 nM camptothecin (+CPT) or 200 nM etoposide (+ETP) for 1 hour and subjected to EdU incorporation for 4 minutes before fixation. **c**. Examples of short and long filaments in EPT-treated cells from Figure 2A,B. **d**. Labeling scheme for the iPOND experiment in Fig 2b and 4a. HEK293T cells were labeled with 10 µM EdU for 10 minutes (UNT) followed by a chase into 10 µM thymidine (Thy) for 50 minutes. In the CPT sample, cells were treated with 100 nM CPT for 60 minutes and EdU was added in the last 10 minutes. Cells were optionally treated with 100 nM LatB 10 minutes prior to CPT treatment (LatB+CPT) and maintained during the genotoxic treatment. **e**. MSD analysis of PCNA foci after indicated treatment (N=1918 foci for UNT and N=2105 foci for +ETP). n=9 cells for the UNT and n=8 cells for +ETP from 3 or more independent experiments. Data are mean ± S.E.M. Statistical analysis: numerical p-values for the indicated comparisons were calculated with extra sum-of-squares *F*-test, nonlinear regression for curve fitting. Δ*t*, time intervals (intervals were 20 s each). Scale bars=5 μm.

**Extended Data Figure 3.**
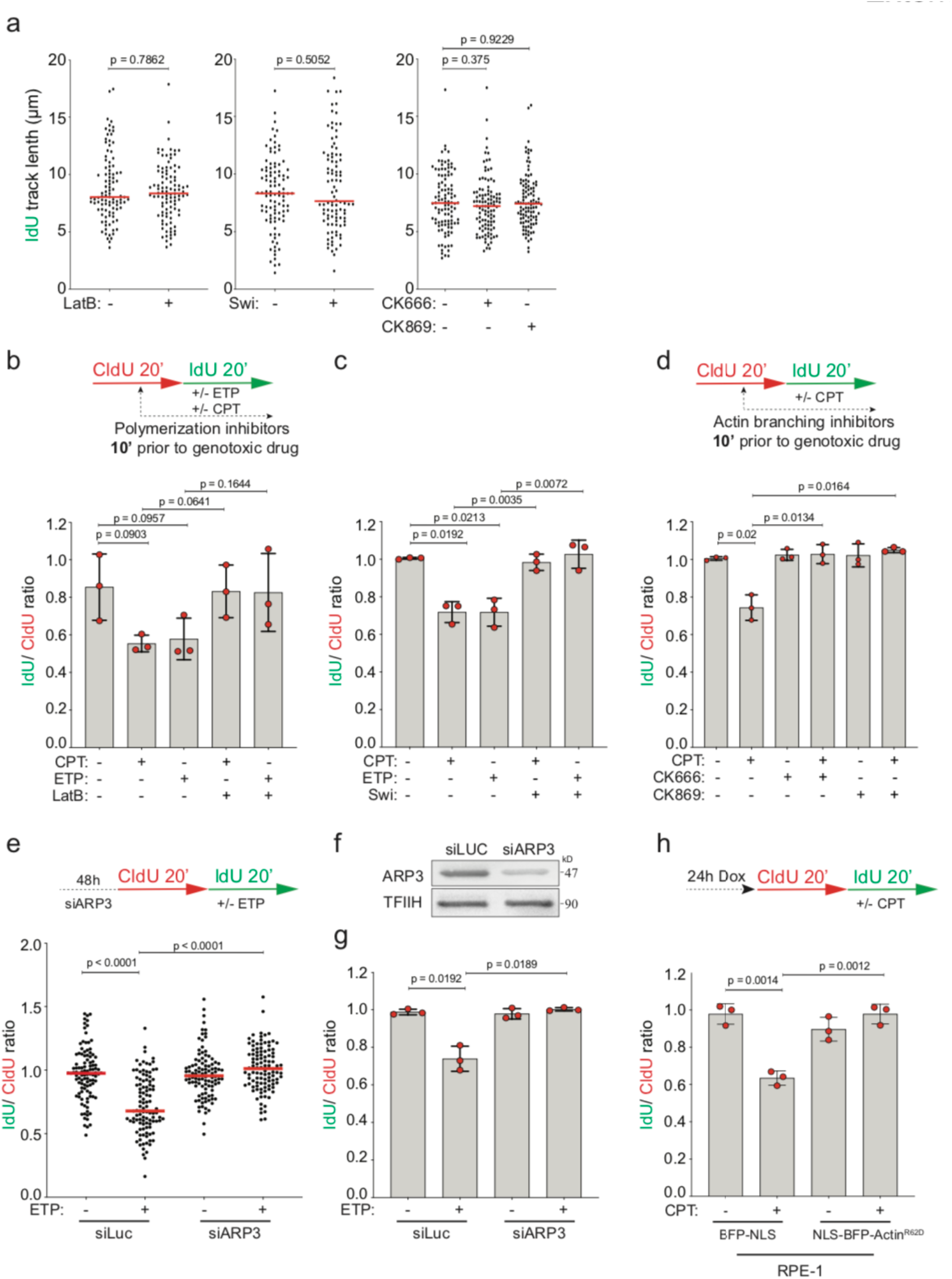
Related to Fig. 3. **a**. DNA fiber analysis of U2OS cells. IdU track length (μm) is plotted as a readout of fork progression. A minimum of 100 forks (indicated as black dots) was scored in three independent experiments yielding similar results. The median values are indicated by horizontal red lines. Statistical analysis: numerical P-values for the indicated comparisons were calculated with Mann–Whitney test. **b-d**. Medians of three biological replicates of fibers in Fig. 3B-D are represented as red dots. The height of the bar represents the mean. 100nM LatB, Swi, CK666 or CK869 were added 10 minutes prior to 100 nM CPT or 20 nM ETP and retained during the IdU labelling. **e**. Top: schematic of the CldU/IdU pulse-labeling protocol used to evaluate fork progression upon 20 nM ETP. U2OS cells were transfected with siARP3 48 hours before CldU and IdU labeling. Bottom: the IdU/CIdU ratio is plotted as a readout of fork progression. Statistical analysis: numerical P-values for the indicated comparisons were calculated with Mann–Whitney test. Medians of three biological fibers replicates are represented as red dots in **g**. The height of the bar represents the mean. **f**. ARP3 levels assessed by western blot. TFIIH, loading control. The experiment was performed three times yielding similar results. h. Top: schematic of the CldU/IdU pulse-labeling protocol used to evaluate fork progression upon 100 nM CPT. Doxycycline (Dox) was added 24 hours before the CldU/IdU pulse-labeling protocol. Bottom: medians of three biological replicates of fibers from Fig.3F are represented as red dots. The height of the bar represents the mean. **b-d, g-h**. A minimum of 100 forks was scored for each replicate. Statistical analysis: numerical P-values for the indicated comparisons were calculated with two tailed Welch’s test.

**Extended Data Figure 4.**
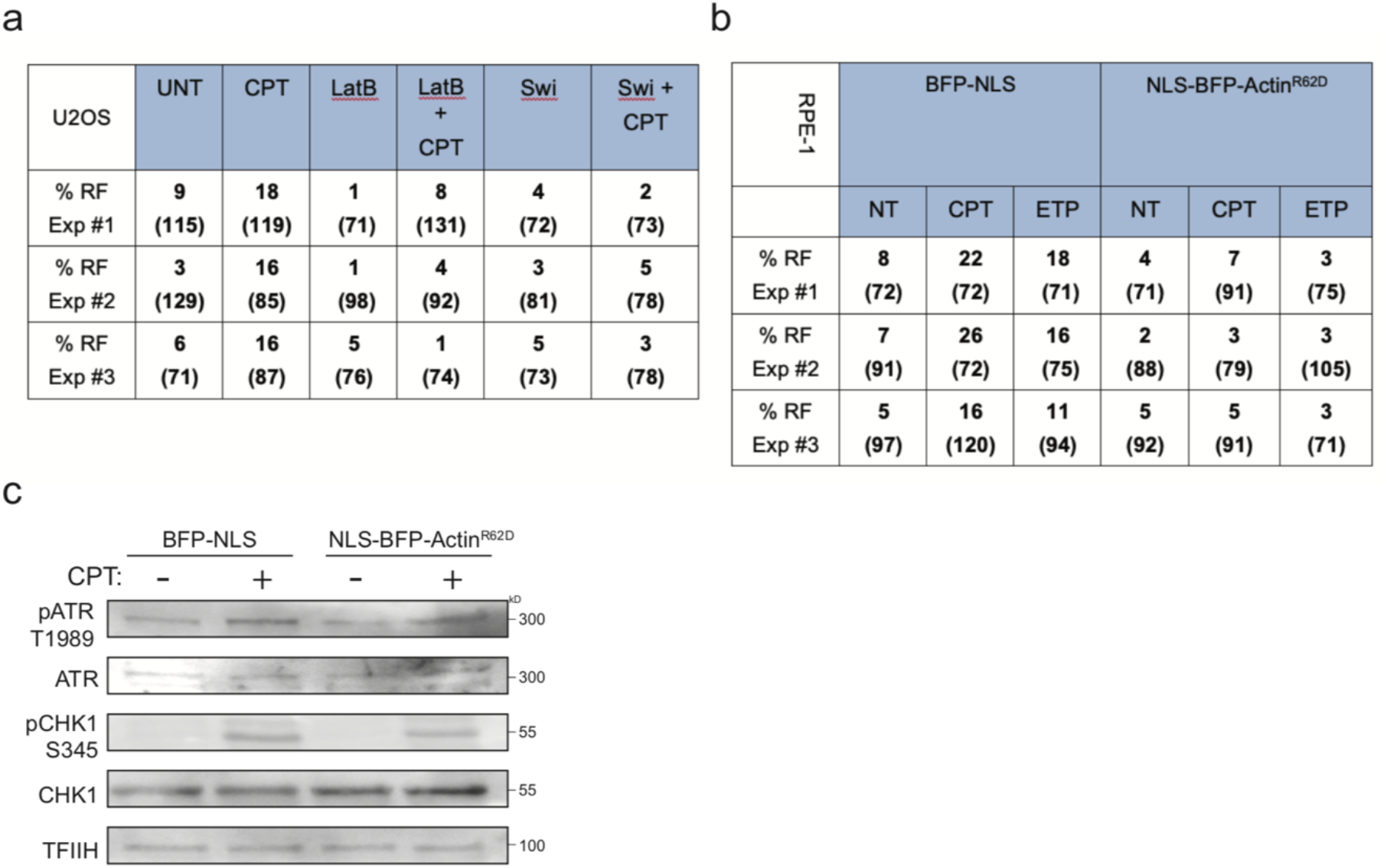
Related to Fig. 4. **a-b**. Electron microscopy summary data. Frequency of observed reversed forks (% RF) in three independent experiments related to Figure 4e-f. Number of analyzed molecules is indicated in brackets. c. Western blot analysis of whole-cell extracts from RPE-1 cells stably expressing doxycycline inducible BFP-NLS or NLS-BFP-ActinR62D. Doxycycline (Dox) was added 24 hours before lysing the cells. BFP-NLS or NLS-BFP-ActinR62D expressing cells were optionally treated for 1 hour with 100 nM CPT. The experiment was performed three times yielding similar results.

**Extended Data Figure 5.**
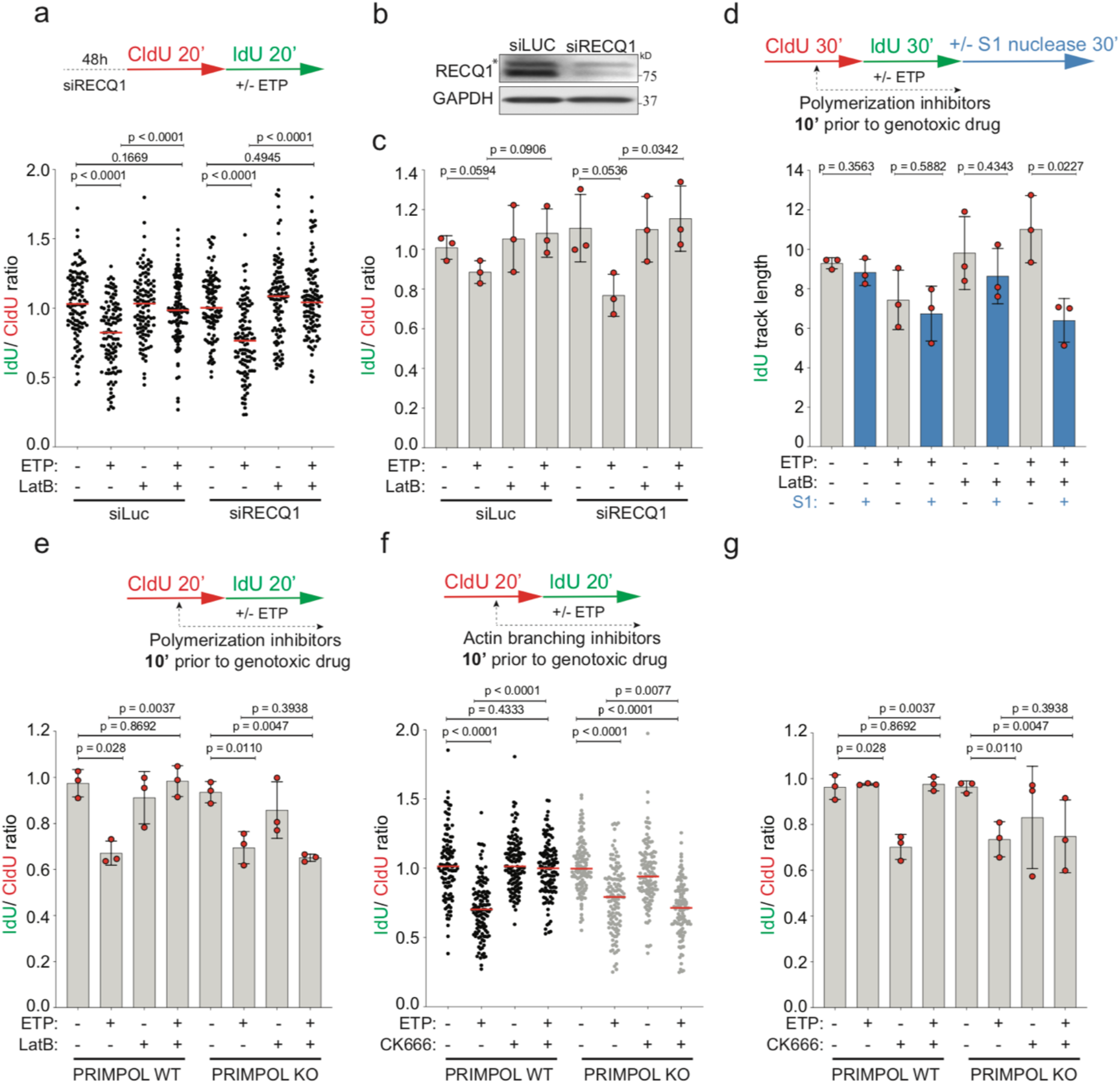
Related to Fig. 5. **a**. Top: schematic of the CldU/IdU pulse-labeling protocol used to evaluate fork progression upon 20 nM ETP. U2OS cells were transfected with siRECQ1 48 hours before CldU and IdU labeling. Bottom: the IdU/CIdU ratio is plotted as a readout of fork progression. A minimum of 100 forks (indicated as black dots) was scored in three independent experiments yielding similar results. The median values are indicated by horizontal red lines. Statistical analysis: numerical p-values for the indicated comparisons were calculated with Mann–Whitney test. Medians of three biological fibers replicates are represented as red dots in **c**. The height of the bar represents the mean. **b**. RECQ1 levels assessed by western blot. GAPDH, loading control; *, unspecific band. The experiment was performed two times yielding similar results. **d-e, g**. Medians of three biological replicates of fibers in Fig. 5b-e and Extended Data Fig. 5fF are represented as red dots. The height of the bar represents the mean. **b-e, g**. A minimum of 100 forks was scored for each replicate. Statistical analysis: numerical p-values for the indicated comparisons were calculated with two tailed Welch’s test. **f**. DNA fiber analysis of U2OS PRIMPOL WT and KO cells. Top: schematic of the CldU/IdU pulse-labeling protocol used to evaluate fork progression upon 20 nM ETP. 100 nM CK666 was added 10 minutes prior to ETP and retained during the IdU labelling. Bottom: the IdU/CIdU ratio is plotted as a readout of fork progression. A minimum of 100 forks was scored in three independent experiments yielding similar results. Statistical analysis: numerical P-values for the indicated comparisons displayed in the figure and calculated with Mann–Whitney test.

**Extended Data Movie 1**, corresponding to Fig. 1b and Extended Data Fig. 1a. U2OS cells stably expressing nAC-GFP were stimulated by 750 nM A23187 to visualize nuclear F-actin polymerization.

**Extended Data Movie 2**, corresponding to Fig. 1b and Extended Data Fig. 1b. Transient formation of nuclear F-actin after cell division in U2OS cells stably expressing nAC-GFP.

**Extended Data Movie 3**, corresponding to Fig. 1d. Distinct and transient nuclear actin filaments are detected in replicating U2OS cells stably expressing nAC-GFP.

## Methods

### Key materials

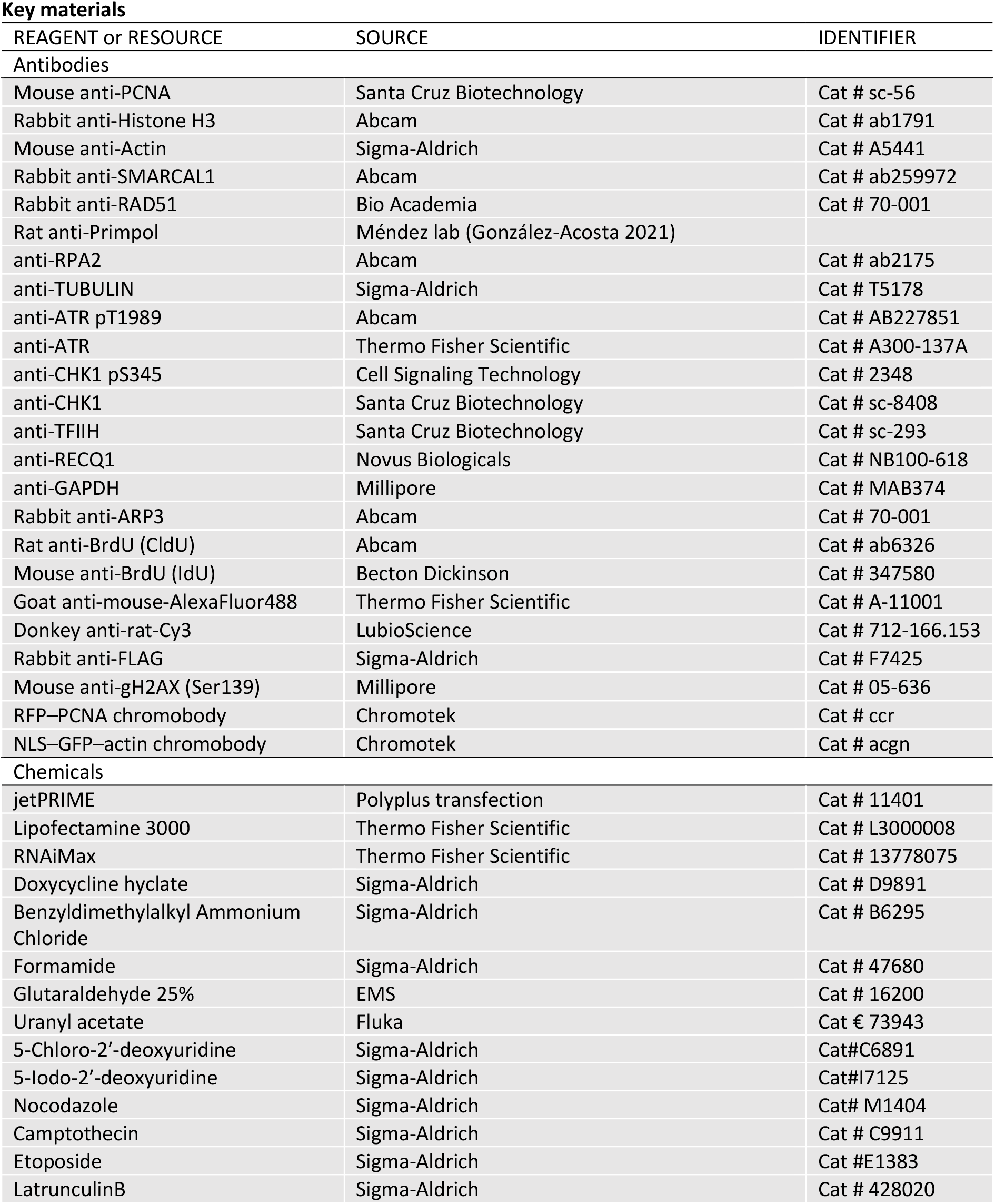
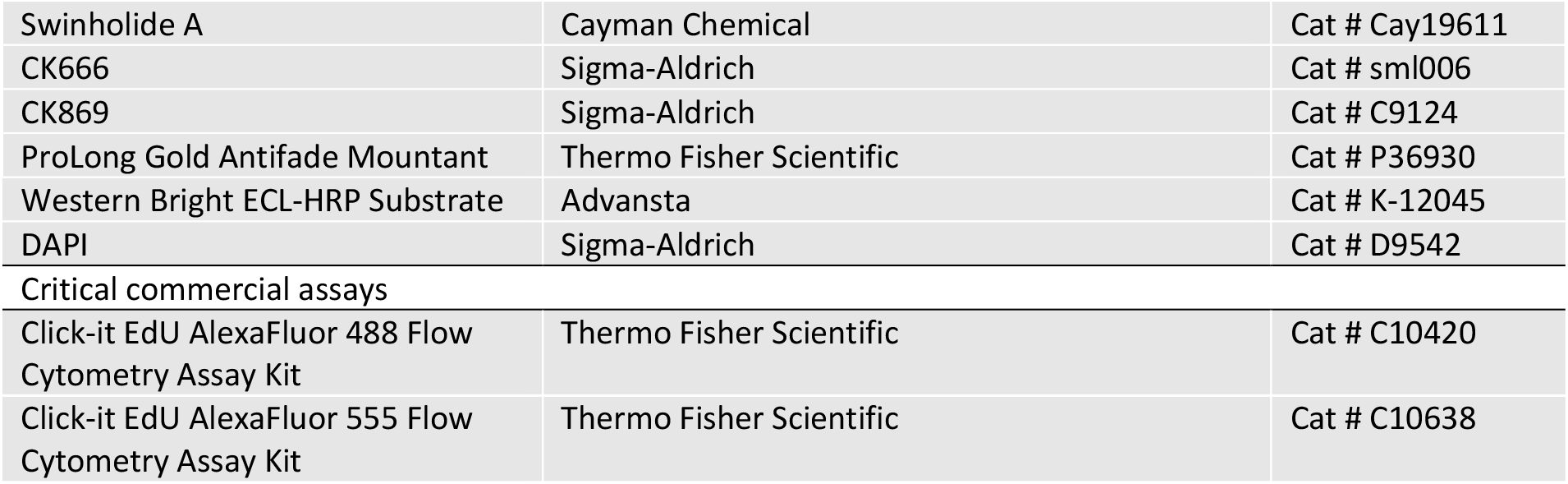

### Cell lines and plasmids

Human osteosarcoma U2OS cells, retinal pigment epithelium RPE-1 cells and HEK293T cells were cultured in DMEM (41966-029, Life Technologies) supplemented with 10% fetal bovine serum (FBS, GIBCO), 100 U/ml penicillin and 100 mg/ml streptomycin in an atmosphere containing 6% CO_2_ at 37°C.

PRIMPOL KO and isogenic U2OS cells were kindly provided by Dr. Juan Méndez.

Nuclear-actin-chromobody (nAC-GFP) stable U2OS cells and stable doxycycline-inducible BFP-NLS or NLS-BFP-Actin^R62D^ cells were kindly provided by Dr. Robert Grosse.

pEF-Flag-NLS-Actin-WT and R62D plasmids were kindly provided by Dr. Guido Posern^44^.

### RNAi experiments

For RNAi experiments, U2OS cells were transfected with the indicated siRNAs for 48 hours: siLuc (5′-CGUACGCGGAAUACUUCGAUUdTdT-3′) and siRECQ1 (5′-UUACCAGUUACCAGCAUUAdTdT-3′); using jetPRIME (Polyplus transfection) according to manufacturer’s instruction. siLuc (5′-CGUACGCGGAAUACUUCGAUUdTdT-3′) and siARP3 (SMARTpool siRNA L-012077-00-0010 (Dharmacon)) were transfected using RNAiMax (Thermo Fisher Scientific) according to manufacturer’s instruction.

### Protein extraction and western blotting

Extracts from all cell lines were prepared in Laemmli sample buffer (4% SDS, 20% glycerol, and 120 mM Tris-HCl, pH 6.8). Equal amounts of protein (30–50 μg) were loaded onto 4%–20% Mini-PROTEAN TGX Precast Protein Gels (BioRad). Proteins were separated by electrophoresis at 16 mA followed by transferring the proteins to Immobilon-P membranes (Thermo Fisher Scientific) for 1 hour at 350 mA (4°C) in transfer buffer (25 mM Tris and 192 mM glycine) containing 10% methanol. Before addition of primary antibodies, membranes were blocked in 5% milk in 0.1% TBST (1 × TBS supplemented with 0.1% Tween 20) for 1 hour and incubated in 3% BSA with primary antibodies overnight at 4°C. Membranes were probed for ADD PRIMARY ANTIBODIES. Secondary antibodies were added for 1 hour at room temperature (in blocking solution). Membranes were washed three times with 0.1% TBST, 10 min each, after primary and secondary antibody incubations and detected with ECL detection reagent (GE healthcare).

### Replication fork progression by DNA fiber analysis

All cell lines subjected to this analysis were grown asynchronously and labeled with 30 μM of the thymidine analog chlorodeoxyuridine (CldU; Sigma-Aldrich) for 20 minutes, they were then washed three times with warm PBS and subsequently exposed to 250 μM of 5-iodo-2′-deoxyuridine (IdU) for 20 minutes alone or in combination with mild doses of genotoxic treatments (100 nM CPT, 20 nM ETP). To evaluate the impact of actin polymerization on replication fork progression, actin polymerization inhibitors were added 10 minutes before the the genotoxic treatment and retained during the IdU labeling. In the RPE-1 cells, 1 μM/mL Dox was added 24 hours before the CldU/IdU pulse labeling. All cells were collected by standard trypsinization and resuspended in cold PBS at 3.5 × 10^5^ cells/mL. 3 μL of this cell suspension were then mixed with 7 μL of lysis buffer (200 mM Tris-HCl, pH 7.5, 50 mM EDTA, and 0.5% [w/vol] SDS) on a glass slide. After an incubation of 9 minutes at RT, the slides were tilted at a 45° angle to stretch the DNA fibers onto the slide. The resulting DNA spreads were air-dried, fixed in 3:1 methanol/acetic acid, and stored at 4°C overnight. The DNA fibers were denatured by incubating them in 2.5 M HCl for 1 hour at RT, washed five times with PBS and blocked with 2% BSA in PBST (PBS and Tween 20) for 40 minutes at RT. The newly replicated CldU and IdU tracks were stained for 2.5 hours at RT using two different anti-BrdU antibodies recognizing CldU (Abcam, ab6326, 1:500) and IdU (Becton Dickinson, 347580, 1:100), respectively. After washing five times with PBST (PBS and Tween 20) the slides were stained with Anti-mouse Alexa 488 (Invitrogen, A-11001, 1:300) and anti-rat Cy3 (Immuno Research, 712-166-1530, 1:150) secondary antibodies for 1 hour at RT in the dark. The slides were mounted in 30 μL Prolong Gold antifade reagent (Invitrogen). For S1 nuclease experiments, cells were treated as described above, labeled with CldU for 30 min, with IdU for 30 min, and incubated with S1 nuclease for 30 min as described^10^.

Microscopy was done using a Leica DM6 B microscope (HCX PL APO 63x objective). To assess fork progression the IdU/CldU ratio or IdU track lengths of at least 100 fibers per sample were measured using the line tool in ImageJ64 software. Statistical analysis was carried out using GraphPad Prism 7.

### Isolation of proteins on nascent DNA or iPOND

iPOND was essentially performed as originally described (Sirbu et al., 2011, 2012) with minor modifications. HEK293T cells were labeled with 10 µM EdU (Life Technologies) for 10 minutes and treated with the different drugs as indicated in Extended Data Fig. 2A. For the pulse-chase experiments with thymidine, after EdU incorporation, cells were washed with cell culture medium and incubated for 50 minutes in medium supplemented with 10 µM thymidine (Sigma-Aldrich). Cells were cross-linked with 1% formaldehyde for 20 minutes at RT, quenched with 0.125 M glycine for 5 minutes, and washed three times with cold PBS. For the conjugation of EdU with biotin azide, cells were permeabilized with 0.25% Triton X-100/PBS for 30 minutes, washed twice with PBS, and incubated in click reaction buffer (10 mM sodium-l-ascorbate, 20 µM biotin azide [Life Technologies], and 2 mM CuSO4) at RT for 2 hours on a rotator. DMSO was used instead of biotin azide for the “no click control”. Cells were washed twice with PBS, resuspended in lysis buffer (50 mM Tris-HCl, pH 8.0, and 1% SDS) supplemented with protease inhibitors (Sigma-Aldrich), and chromatin was solubilized by sonication in a Bioruptor (Diagenode) at 4°C at the highest setting for 10 min (30 s on and 30 s off cycles). After centrifugation for 10 min at 16,000 RCF, supernatants were diluted with 1:1 PBS (vol/vol) containing protease inhibitors and incubated overnight with streptavidin-agarose beads (EMD Millipore). Beads were washed once with lysis buffer, once with 1 M NaCl, twice with lysis buffer, and once with PBS, and captured proteins were eluted by boiling beads in 2× NuPAGE LDS Sample Buffer (Life Technologies) containing 100 mM DTT for 45 minutes at 95°C. Proteins were resolved by electrophoresis using NuPAGE Novex 4–20% Bis-Tris gels and detected by Western blotting with the indicated antibodies: rabbit anti-Histone H3 (1:10000, abcam; ab1791), mouse anti-PCNA (1:2000, Santa-Cruz; sc-56), mouse anti-Actin (1:1000, Abcam; A5441), rabbit anti-SMARCAL1 (1:1000, abcam; ab2559972), rabbit anti-RAD51 (1:1000, BioAcademia; 70-001).

### Colony forming assay

RPE-1 cells stably expressing BFP-NLS or NLS-BFP-Actin^R62D^ were induced with 1 μM/mL Dox 24 hours before performing the experiment. Cells were then optionally treated with 50 nM or 200 nM CPT for 1 hour, washed, harvested and re-plated in triplicates into 6-well plates at low cell dilutions and subsequently cultured for 10 days to allow formation of colonies. Subsequently, colonies were stained with 0.5% (w/v) crystal violet in 20% ethanol. Plates were imaged using a plate reader and colony forming area quantified using the ColonyArea ImageJ plugin as described^66^.

### Analysis of chromosome spreads

U2OS cells were treated with 100 nM CPT for 2 hours. Where indicated, cells were pretreated with 100 nM LatB 10 minutes prior to CPT and maintained during the genotoxic treatment. RPE-1 cells stably expressing NLS-Actin-WT or NLS-Actihn-R62D were induced with 1 μM/mL Dox 24 hours before performing the experiment. The cells were then treated with 50 nM CPT for 8 hours. The genotoxic agents were removed by washing three times with 1 × PBS and the cells were then released into fresh medium containing 200 ng ml^-1^ nocodazole for 16 hours. Cells were collected and swollen with 75 mM KCl for 20 min at 37 °C. Swollen mitotic cells were collected and fixed with methanol:acetic acid (3:1). The fixing step was repeated two times. Cells were then dropped onto pre-hydrated glass slides and air-dried overnight. The following day, slides were mounted with Vectashield medium containing DAPI. Images were acquired with a microscope (model DMRB; Leica) at 63x magnification equipped with a camera (model DFC360 FX; Leica) and visible chromosome abnormalities per metaphase spread were counted.

### Chromatin fractionation

U2OS cells were treated with 100 nM CPT for 1 hour. Where indicated, cells were pretreated with 100 nM LatB 10 minutes prior to CPT and maintained during the genotoxic treatment. Whole cell extracts were prepared by suspension of cells in Laemmli buffer (50 mM Tris–HCl pH 6.8, 10% glycerol, 3% SDS, 0.006 w/v bromophenol blue and 5% 2-mercaptoethanol) followed by sonication. Biochemical fractionation was performed as described (Mendez & Stillman, 2000). SDS–PAGE, protein transfer to nitrocellulose, and immunoblots were performed using standard protocols.

### Electron microscopic analysis of genomic DNA

Asynchronous and subconfluent U2OS cells were treated with 100 nM CPT for 1 hour. Where indicated, cells were pretreated with 100 nM LatB or Swi 10 minutes prior to CPT and maintained during the genotoxic treatment. Asynchronous and subconfluent RPE-1 cells stably expressing NLS-Actin-WT or NLS-Actihn-R62D were induced with 1 μM/mL Dox 24 hours before collecting the cells. The cells were then treated with 100 nM CPT or 20 nM ETP for 1 hour. Cells were collected, resuspended in ice-cold PBS and crosslinked with 4,5′, 8-trimethylpsoralen (10 μg/ml final concentration), followed by irradiation pulses with UV 365 nm monochromatic light (UV Stratalinker 1800; Agilent Technologies). For DNA extraction (Muzi-Falconi and Brown, 2018), cells were lysed (1.28 M sucrose, 40 mM Tris-HCl [pH 7.5], 20 mM MgCl2, and 4% Triton X-100; Qiagen) and digested (800 mM guanidine–HCl, 30 mM Tris-HCl [pH 8.0], 30 mM EDTA [pH 8.0], 5% Tween-20, and 0.5% Triton X-100) at 50 °C for 2 h in presence of 1 mg/ml proteinase K. The DNA was purified using chloroform/isoamylalcohol (24:1) and precipitated in one volume of isopropanol. Finally, the DNA was washed with 70% EtOH and resuspended in 200 μl TE (Tris-EDTA) buffer. 120 U of restriction enzyme (PvuII high fidelity, New England Biolabs) were used to digest 6 μg of the purified genomic DNA for 5 h at 37C. RNase A (Sigma–Aldrich, R5503) to a final concentration of 250 ug/ml was added for the last 2 h of this incubation. The digested DNA purified using the Thermo Fisher Silica Bead Gel Extraction kit according to manufacturer’s instructions. The Benzyldimethylalkylammonium chloride (BAC) method was used to spread the DNA on the water surface and then load it on carbon-coated 400-mesh nickel grids (G2400N, Plano Gmbh). Subsequently, DNA was coated with platinum using a High Vacuum Evaporator BAF060 (Leica). The grids were imaged automatically using a Talos 120 transmission electron microscope (FEI; LaB6 filament, high tension ≤120 kV) with a bottom-mounted CMOS camera BM-Ceta (4000×4000pixel) and the MAPS software package (Thermo Fisher Scientific, Eindhoven, The Netherlands) as described^51^. Samples were annotated for replication intermediates using the MAPS Viewer software, overlapping images for annotated regions were stitched together using the automated pipeline ForkStitcher^51^ and final images were analyzed using ImageJ. For each experimental condition at least 70 replication fork molecules were analyzed in three different biological replicates.

### Immunofluorescence and imaging of fixed samples

U2OS cells were transiently transfected with FLAG-NLS-WT or FLAG-NLS-R62D-Actin for 24 hours with Lipofectamine 3000 (Thermo Fisher Scientific) according to the manufacturer’s instructions and grown on sterile 12-mm diameter glass coverslip coated with poly-L-lysine, incubated for 4 minutes with 10 μM EdU, washed with 1X PBS and pre-extracted for 10 min with CSK-buffer (10 mM PIPES, 50 mM NaCl, 300 mM Sucrose, 3 mM MgCl_2_, 1 mM EGTA, and 0.5% Triton X-100) on ice, fixed in 4% buffered paraformaldehyde, washed three times with 1X PBS, permeabilized for 10 min at room temperature in 0.3% Triton X-100 (Sigma-Aldrich) in PBS and washed twice in PBS. EdU detection was performed with a Click-iT Plus EdU Alexa-Fluor 555 Imaging Kit according to the manufacturer’s recommendations (Thermo Fisher Scientific) before incubation with primary antibodies. All primary and secondary antibodies were diluted in 3% BSA/PBS. Incubation with primary antibodies was performed at room temperature for 2 hours or overnight at 4°C. Coverslips were washed three times with PBS containing 0.1% Tween-20 (Sigma-Aldrich). Secondary-antibody incubations were performed at room temperature for 45 minutes. After one wash with PBS containing 0.1% Tween-20 and one with PBS, coverslips were incubated for 10 min with PBS containing DAPI (0.5 mg/mL) at room temperature to stain DNA. Following three washing steps in PBS, coverslips were briefly washed with distilled water, dried on 3 mm paper and mounted with Prolong Gold antifade reagent (Invitrogen). Imaging and image processing for fixed cells done in Fig. 2A, C and in Extended fig. 2C has previously been described (add ref 66 See et al 2020 and 67 Caridi et al 2018). Confocal microscopy of a single middle *Z-*stack from fixed S-phase (EdU+) U2OS cells in Extended figure 2A was done using TCS SP5 Leica confocal microscope equipped with the HCX PL APO ×63 oil-immersion objective (NA, 1.4).

### Transient transfection and live cell imaging

The PCNA-chromobody (PCNA-CB-RFP) (Chromotek, ccr) was purchased with a material transfer agreement from Chromotek. It was transiently transfected in nAC-GFP stable U2OS cells with Lipofectamine 3000 (ThermoFisher Scientific) according to the manufacturer’s instructions and 24 hours before imaging. Cells were then seeded on a previously poly-L-lysine coated μ-slides glass-bottom dish (Ibidi). Culture medium was replaced with colourless DMEM (FluoroBrite DMEM, ThermoFisher Scientific) 2 hours before imaging. Imaging was performed using the incubation system for live cell imaging (cellVIVO) at 37 °C and 5% CO2 on a Olympus IXplore SpinSR10 (Olympus Europe SE & Co. KG, Germany) with a Yokogawa CSU-SoRa disk for super resolution imaging and UPLSAPO 100x/1.3 NA oil-immersion objective. The acquisition software CellSens Dimension 3.1 (Olympus) was used. In order to avoid phototoxicity and photobleaching, images were acquired every 20 seconds (imaging frames) for three time intervals of 5 minutes (T1,T2,T3 in the figures) separated by 5 minutes dark intervals. Z series were collected with 0.3 μm intervals over a 12 μm range. 200 nM ETP or H2O (UNT) were added to the cells just before imaging. Mitotic cells were also imaged every 20 seconds until detectable exit from mitosis, whereas those treated with 750 nM A23817 were imaged every minute until clear F-actin dissolution.

### Live cell imaging processing

PCNA foci were tracked in 3D using a semi-automated method and manually corrected to ensure optimal connections between time points as previously described^67,68^. Tracked foci were used to register the cells. After registration, PCNA foci were retracked and focus positional data were extracted in Excel and analyzed in Matlab (MathWorks) to calculate MSDs as previously described^67,68^. Quantification of actin filament length was done using the Measurement Tool in Imaris. When a filament persists for several time points, only the time point with the longest filament was measured. Videos were generated using individual snapshots saved as .tiff files and assembled in Fiji.

### Quantification and image processing af F-actin

Image processing for live and fixed cell data of nuclear F-actin was performed with IMARIS (Bitplane), FIJI (NIH) and Photoshop CC (Adobe). In Fig. 2A-B, cells were scored as positive when nuclear actin filaments were detected and as negative when no nuclear actin filaments were observed. Cells with nuclear actin filaments were further classified into subcategories; short and long filaments or a combination of these. No data were excluded from the analysis.

## Acknowledgments

We are grateful to Jonas A. Schmid, Gabriel Senn and the Center for Microscopy and Image Analysis of the University of Zurich for technical assistance with microscopy and imaging analysis. We thank M. Altmeyer and all members of the Lopes lab for useful discussions and suggestions on the manuscript. Work in the Lopes lab was supported by the SNF Project grant 310030_189206 and a grant from the Novartis Foundation. M.D.P. was supported by an AIRC “Fellowship for Abroad” and by the Forschungskredit of the University of Zurich. Work in the Chiolo lab was supported by R01GM117376 and NSF Career 1751197.

